# Microbiota encoded fatty-acid metabolism expands tuft cells to protect tissues homeostasis during *Clostridioides difficile* infection in the large intestine

**DOI:** 10.1101/2024.01.29.574039

**Authors:** Tasia D. Kellogg, Simona Ceglia, Benedikt M. Mortzfeld, Abigail L. Zeamer, Sage E. Foley, Doyle V. Ward, Shakti K. Bhattarai, Beth A. McCormick, Andrea Reboldi, Vanni Bucci

## Abstract

Metabolic byproducts of the intestinal microbiota are crucial in maintaining host immune tone and shaping inter-species ecological dynamics. Among these metabolites, succinate is a driver of tuft cell (TC) differentiation and consequent type 2 immunity-dependent protection against invading parasites in the small intestine. Succinate is also a growth enhancer of the nosocomial pathogen *Clostridioides difficile* in the large intestine. To date, no research has shown the role of succinate in modulating TC dynamics in the large intestine, or the relevance of this immune pathway to *C. difficile* pathophysiology. Here we reveal the existence of a three-way circuit between commensal microbes, *C. difficile* and host epithelial cells which centers around succinate. Through selective microbiota depletion experiments we demonstrate higher levels of type 2 cytokines leading to expansion of TCs in the colon. We then demonstrate the causal role of the microbiome in modulating colonic TC abundance and subsequent type 2 cytokine induction using rational supplementation experiments with fecal transplants and microbial consortia of succinate-producing bacteria. We show that administration of a succinate-deficient *Bacteroides thetaiotaomicron* knockout (Δfrd) significantly reduces the enhanced type 2 immunity in mono-colonized mice. Finally, we demonstrate that mice prophylactically administered with the consortium of succinate-producing bacteria show reduced *C. difficile*-induced morbidity and mortality compared to mice administered with heat-killed bacteria or the vehicle. This effect is reduced in a partial tuft cell knockout mouse, *Pou2f3*^+/-^, and nullified in the tuft cell knockout mouse, *Pou2f3*^-/-^, confirming that the observed protection occurs *via* the TC pathway. Succinate is an intermediary metabolite of the production of short-chain fatty acids, and its concentration often increases during dysbiosis. The first barrier to enteric pathogens alike is the intestinal epithelial barrier, and host maintenance and strengthening of barrier integrity is vital to homeostasis. Considering our data, we propose that activation of TC by the microbiota-produced succinate in the colon is a mechanism evolved by the host to counterbalance microbiome-derived cues that facilitate invasion by intestinal pathogens.

## Introduction

Microbiota-produced metabolites have a crucial role in modulating local and peripheral immune signatures (Blander *et al*., 2017; McCarville *et al*., 2020) with three prominent examples that include short chain fatty acids (SCFAs) (Arpaia *et al*., 2013; Atarashi *et al*., 2013; Atarashi *et al*., 2011; Foley *et al*., 2021; Schulthess *et al*., 2019; Tanoue *et al*., 2016), indoles (Aoki *et al*., 2018; Goettel *et al*., 2016) and secondary bile acids (Foley Sage *et al*., 2022; Ridlon *et al*., 2014).

Previous research demonstrates that succinate produced by parasites in the small intestine (SI) is sensed by the taste-chemosensory epithelial tuft cells (TCs) (Chen *et al*., 2022; Howitt *et al*., 2016; O’Leary *et al*., 2019) and initiates a type 2 immune signaling cascade leading to parasite expulsion. (Loke and Cadwell, 2018; Luo *et al*., 2019; Miller *et al*., 2018; Nadjsombati *et al*., 2018; Schneider *et al*., 2018a; von Moltke *et al*., 2016). TCs secrete IL-25 which acts on type 2 innate lymphoid cells (ILC2s) to produce the type 2 cytokines IL-4, IL-5, and IL-13 (Loke and Cadwell, 2018; Luo *et al*., 2019; Miller *et al*., 2018; Nadjsombati *et al*., 2018; Schneider *et al*., 2018a; von Moltke *et al*., 2016). These type 2 cytokines act synergistically to cause hyperplasia of TCs and mucus-producing goblet cells (Schneider *et al*., 2018b), increased SI length, increased contractility of the smooth muscle within the intestine, and recruitment of eosinophils to the epithelial barrier (Howitt *et al*., 2016). The physiological outcome of this process, usually referred to as the “weep and sweep” response (WSR), is the expulsion of the parasites *via* mucus production (weep) and increased motility (sweep) (Howitt *et al*., 2016; von Moltke *et al*., 2016).

To date, most of the literature that details the function of TCs in the GI tract focuses on the small intestine and how these small intestinal TCs interact with, respond to intestinal parasites and their substrates, and initiate protection against them. Despite the interest in TCs in the GI tract and beyond, there is limited information about colonic TCs. TCs have been previously identified in the colon, and shown to possess the succinate receptor, SCNR1 (Lei *et al*., 2018). While some literature has demonstrated a potential impact of bacterial members of the microbiota on TCs, these papers have been limited to the small intestine and only relied on systemic antibiotic manipulation of the microbiome (Banerjee *et al*., 2020). Indeed, despite the fact that the microbiota predominantly occupies the colon compared to the SI by multiple orders of magnitude (Kennedy and Chang, 2020), there is no report about communication between TCs and commensal microbes in the colon.

In the colon, succinate accumulation often occurs when the microbiome is perturbed (Tulstrup *et al*., 2015) and it has been considered a biomarker of inflammation since higher succinate concentrations are observed in the serum and feces of IBD patients compared to healthy controls (Fremder *et al*., 2021). Succinate is the most prevalent biochemical route to propionate production by primary fermenters including members of the *Bacteroides* and *Prevotella* genera (Ikeyama *et al*., 2020). It is also a major cross-feeding metabolite (Fernández-Veledo and Vendrell, 2019), was shown to enhance the *in vivo* growth of *Clostridioides difficile* (Ferreyra *et al*., 2014), and acts as an environmental signal to regulate *Salmonella’s* virulence as well as host invasion programs (Spiga *et al*., 2017).

Type 2 immunity, a hallmark response to helminths and allergens, also mediates tissue regeneration in many muco-cutaneous barriers including the colon (Akdis *et al*., 2020; Cox *et al*., 2021; Gieseck *et al*., 2018). In mice, intraperitoneal administration of recombinant IL-25 induced eosinophil-mediated barrier protection against *Clostridioides difficile* morbidity and mortality with no effect on *C. difficile* intestinal levels (Buonomo *et al*., 2016). Similar findings were obtained in humans with lower IL-25 concentrations found in colonic biopsies of *C. difficile*-infected patients compared to healthy controls (Buonomo *et al*., 2016). Finally, TCs and TC-derived IL-25 were also protective in a mouse model of DSS-induced colitis (Qu *et al*., 2015), and patients with inflammatory bowel disease (IBD) display fewer IL-25 expressing cells in their intestinal mucosa, with IL-25 levels being lower during active disease compared to remission (Su *et al*., 2013). Thus, TCs response is critical for maintain homeostasis of the colonic tissue in many inflammatory and auto-inflammatory conditions, suggesting starkly distinct TCs function according to anatomy (i.e., colon vs. the small intestine), possibly imprinted by host and environmental cues.

Here we hypothesize that microbially-produced succinate is a metabolite at the center of a three-way circuit that includes the microbiome, *C. difficile,* and host epithelial cells in the colon. Specifically, we propose that colonic TCs expansion in response to the accumulation of microbiota-produced succinate acts as a protective mechanism by which the host resolves *C. difficile*-caused intestinal distress. Through a combination of selective microbiome disruption and supplementation experiments with antibiotics, fecal matter transplantation (FMT), and administration of defined consortia of genetically competent and altered bacteria, we demonstrate that the expansion of TCs in the colon and production of type 2 cytokines crucially depends on the succinate produced by colonic bacteria. We further demonstrate that the administration of succinate-producing bacteria leads to protection against *C. difficile-*induced morbidity and mortality *via* this TC-activated pathway. Using *Pou2f3*^-/-^ mice that lack the transcription factor *Pou2f3* crucial for the differentiation of DCLK1+ TCs, we confirm that protection from *C. difficile* pathogenesis is mediated through colonic tuft cell expansion.

## Results

### Vancomycin treatment causes an increase in colonic IL-25, IL-13, IL-5, and TCS number

The antibiotic vancomycin, which targets Gram-positive bacteria including many succinate-consuming commensal *Clostridia* (Isaac *et al*., 2017), has been shown to preferentially and specifically increase IL-25 production in the colon (Tulstrup *et al*. 2015, Li *et al*. 2019). Consequently, we hypothesized that vancomycin would also promote TCs expansion and thereby, in addition to IL-25, lead to IL-13 and IL-5 concentrations. To test this, we compared tissues from mice selectively administered with antibiotics, either vancomycin, metronidazole, an antibiotic cocktail (“AVNM,” containing ampicillin, vancomycin, neomycin, and metronidazole), or sterile phosphate-buffered saline (PBS) for seven days by oral gavage. We assessed the differences in IL-25, IL-13, and IL-5 concentrations by first using enzyme-linked immunosorbent assays (ELISA) (see Methods). We found higher colonic IL-25 protein concentrations in vancomycin-treated mice compared to untreated (p=0.001), AVNM-treated (p=0.001) and metronidazole-treated (p=0.001) mice (**Fig. 1A**). Levels of IL-5 and IL-13 proteins were also significantly elevated in the colon of vancomycin-treated mice compared to untreated, metronidazole-treated, or AVNM-treated mice (p<0.05) (**Fig. S1**). Interestingly, no increase in IL-25, IL-13 or IL-5 was observed in the cecum or the ileum (**Fig. S2**). Reverse transcription-quantitative polymerase chain reaction (RT-qPCR) performed on a subset of samples confirmed the IL-25 results. Colonic IL-25 expression was significantly increased in vancomycin-treated mice (p-value ANOVA with Tukey post-hoc = 0.001) compared to mice that received PBS (e.g., untreated), or metronidazole (p = 0.01) (**Fig. 1**). We then evaluated TCs levels by assessing the ratio between of DCLK1+ expressing epithelial cells relative to the number of epithelial (EPCAM+) cells *via* flow cytometry (von Moltke *et al*., 2016) (**Fig. 1C**, **Fig. S3**) (See Methods). Mice treated with vancomycin showed significantly higher proportions of TCs compared to mice receiving PBS (p = 0.005), and a marginally significant increase compared to metronidazole (**Fig. 1C**). The observed phenotype was independent of intestinal colonization by fungi, which are potential succinate producers (Begum *et al*., 2022). This was confirmed by treatment with the antifungal amphotericin B (See Methods) which resulted in no change to the enhanced IL-25, IL-13, and IL-5 protein concentrations in vancomycin-treated animals (**Fig. S4**). Our data show that vancomycin administration results in the expansion of TCs, and an increase in the levels of IL-25, IL-13, and IL-5 in the proximal colon.

**Figure 1:**
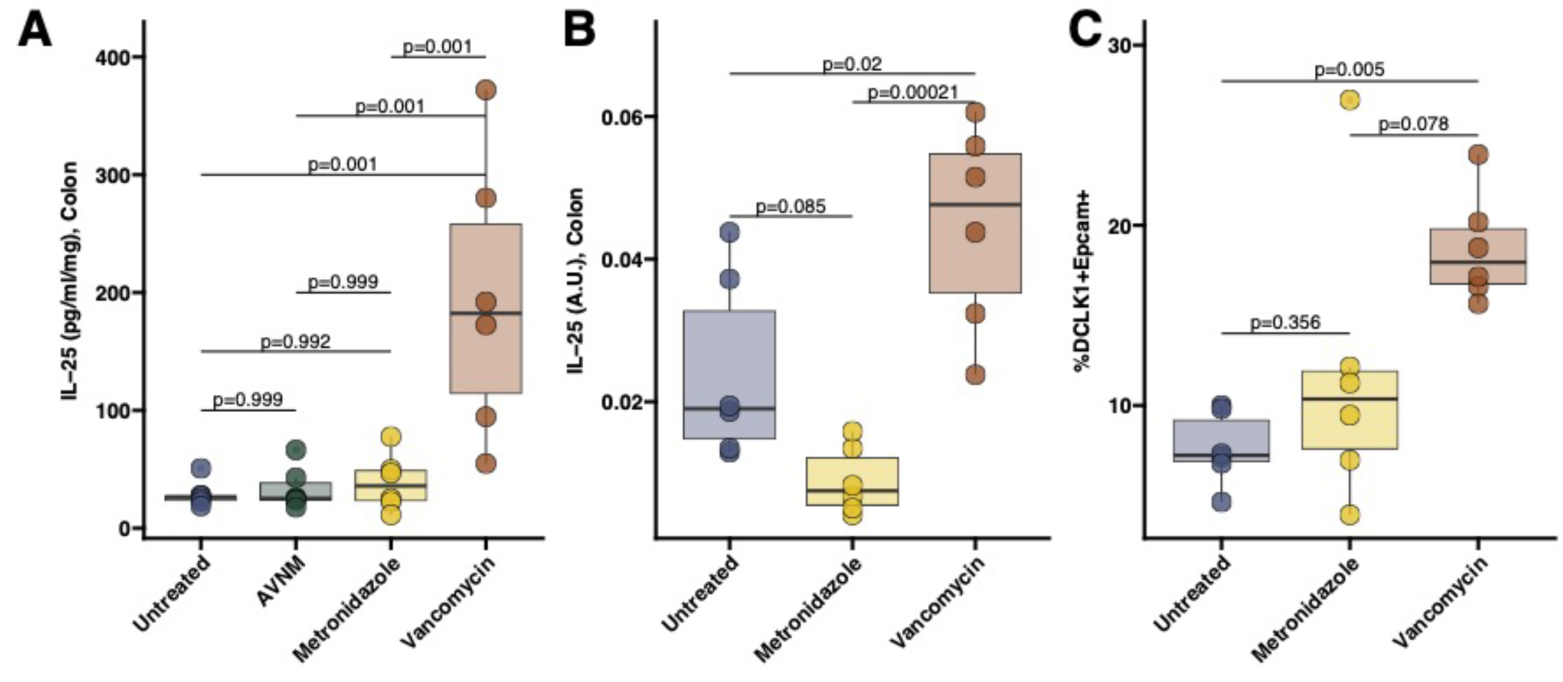
Vancomycin administration expands tuft cells and IL-25 in the proximal colon. **A.** Following the approach from (Buonomo *et al*., 2016) IL-25 was measured by ELISA in the proximal colon of C57BL/6 wild-type mice treated with various antibiotics *via* oral gavage. **B.** IL-25 mRNA expression was measured by RT-qPCR in the proximal colon of C57BL/6 wild-type mice treated with different antibiotics. **C.** Percentage of DCLK1+ EPCAM+ CD45-cells compared to total cells in C57BL/6 wild-type mice treated with various antibiotics or untreated.

### Microbiome reconstitution with FMT enriched in succinate-producing bacteria increases IL-25, IL-13, and IL-5 in the colon but not in the cecum or ileum

To demonstrate a causal role for the microbiome in inducing colonic IL-25, IL-13, and IL-5, we performed FMT experiments as previously described (Ubeda *et al*., 2013). Mice were either pre-treated with AVNM ad-libitum in drinking water for 7 days or left untreated (i.e., sterile water). Following pre-treatment, mice were orally gavaged with a stool fraction from mice previously treated with vancomycin for one week and weaned off the antibiotic prior to donor stool collection, or from mice that remained on sterile water (as control). We observed a significant increase in the colonic levels of all three cytokines in AVNM-pretreated mice receiving vancomycin-treated stool compared to untreated stool (FDR adjusted p-value ANOVA with Tukey post-hoc < 0.05) (**Fig. 2A**). No difference was observed in mice that were not AVNM-pretreated (**Fig. 2A**). No significant difference in IL-25 was observed in the cecum or the ileum between mice receiving vancomycin-treated stool or untreated stool irrespective of the pre-treatment background (**Fig. 2B**), suggesting that this microbiome-dependent induction colocalizes with the intestinal site of engraftment in the colon (Smillie *et al*., 2018).

**Figure 2:**
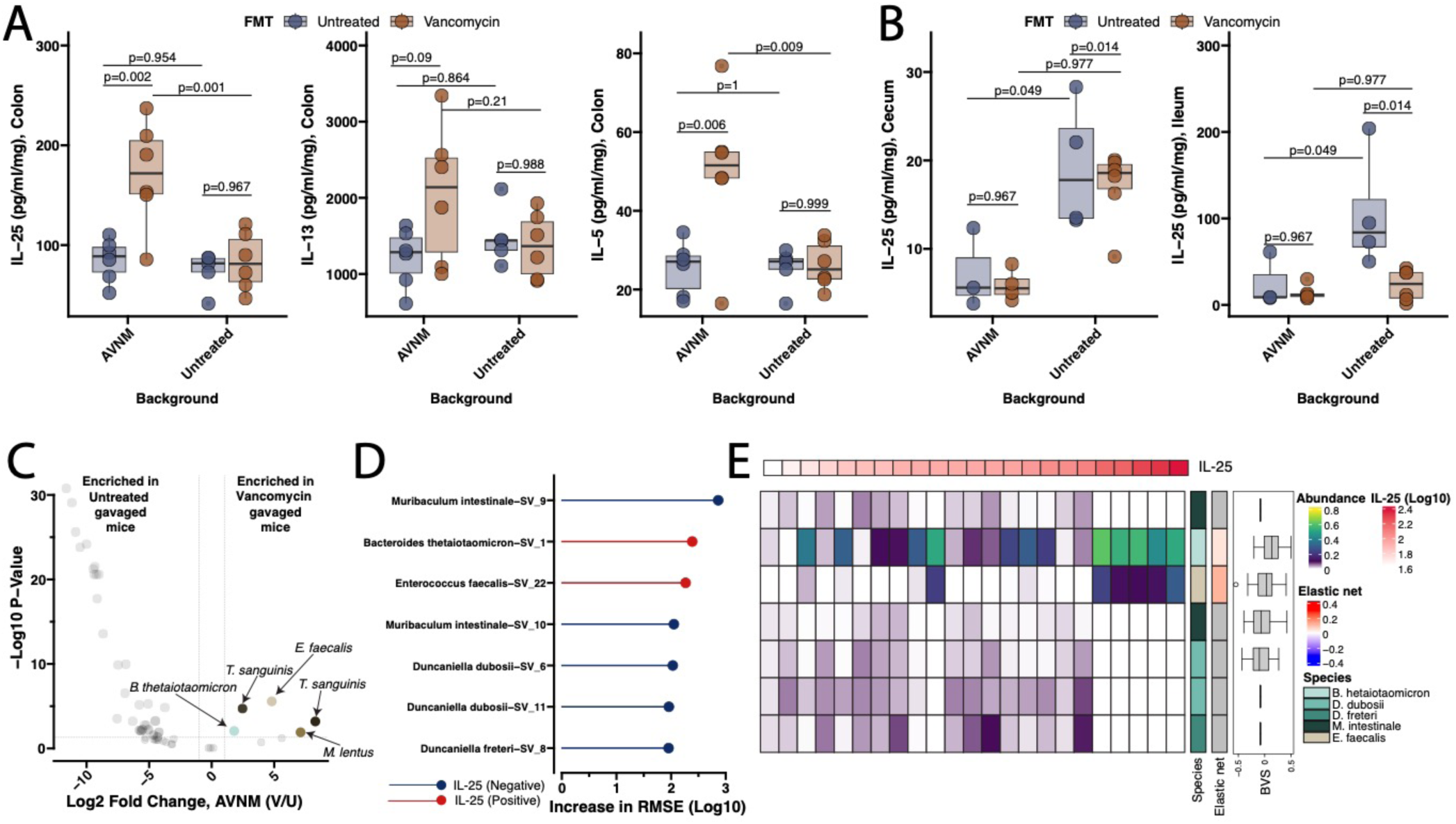
Fecal microbiota transplantation experiments demonstrate the causal role of microbiome in the induction of Type 2 cytokines in the proximal colon, which correlates with the enrichment of *Bacteroides thetaiotaomicron* in the microbiome. **A.** IL-25, IL-5 and IL-13 levels were measured in the proximal colon of C57BL/6 wild-type mice (see Methods) receiving fecal matter transplants (FMTs) from mice treated with vancomycin or left untreated. Before the FMT mice were either left untreated or were pretreated with an antibiotic combination of ampicillin, vancomycin, neomycin, and metronidazole (AVNM) to clear the resident microbiota (Background). **B.** IL-25 levels were assessed in the cecum and ileum. **C.** Volcano plots were obtained after running differential analysis using DeSeq2 on the fecal microbiome sequencing samples from mice pretreated with AVNM and receiving FMT from either a vancomycin-treated or an untreated donor. Two known succinate-producing species (*B. thetaiotaomicron* and *E. faecalis*) are enriched in vancomycin-treated FMT recipients. **D.** Permutated importance analysis along with accumulated local effects calculation was performed on the results of the Random Forest Regression (RFR) modeling to predict colonic IL-25 as a function of microbiome species abundance. Sequence variants belonging to *B. thetaiotaomicron* and *E. faecalis* were determined to positively predict IL-25 concentrations. **E**. Significance and directionality inferred using the RFR model were confirmed running Elastic Net Regression modeling and Bayesian Variable Selection Linear Regression. ASV belonging to *B. thetaiotaomicron* and *E. faecalis* are found to significantly predict accumulation of IL-25 in the colon. Protein concentration in the lysate (pg/mL) was normalized by the total protein mass generated in the sample (in mg)

To determine differences in the microbiome of mice that differentially responded to FMTs, we performed 16S rRNA sequencing of fecal samples from all mice obtained 24 hours after the last FMT. No significant differentially-abundant species were observed post-transplant in mice receiving FMT from vancomycin-treated mice or untreated mice if they did not receive a pre-treatment with AVNM (FDR adjusted p-value > 0.05 using DeSeq2, see (Love *et al*., 2014)). This is expected, as either perturbed or naturally dysbiotic microbiotas are needed in order to facilitate colonization by FMT-derived bacteria (Suez *et al*., 2018). For the mice that received AVNM pretreatment, amplicon sequencing variants (ASVs) belonging to succinate-producing species of *Bacteroides thetaiotaomicron* and *Enteroccocus faecalis* (Catlett *et al*., 2020; Kim *et al*., 2016) were observed to be significantly increased in relative abundance in mice receiving stool from vancomycin-treated relative to untreated mice (**Fig. 2C**) (FDR adjusted p-value <0.05 using DeSeq2, see (Love *et al*., 2014)). To identify which bacterial species were associated with the observed host induction in the AVNM-pretreated mice we took advantage of random forest regression (RFR) modeling as we have previously described (Wipperman *et al*., 2021). Specifically, we used RFR to predict the concentration of IL-25 in every sample as a function of the abundances of all detected ASVs in these mice (See Methods). ASVs mapping to *B. thetaiotaomicron* and *E. faecalis* were found to be the only two bacteria (**Fig. 2D**) whose increase in abundance predicts a significant increase in colonic IL-25 levels. Positive significant associations between the abundance of *B. thetaiotaomicron* and *E. faecalis* ASVs, and colonic IL-25 levels were then corroborated by running both Elastic Net as well as Bayesian Variable Selection linear regression modeling on this data (**Fig. 2E**). Taken together, these data show that microbiome reconstitution with stool fractions leading to the enrichment in succinate-producing bacteria coincides with the upregulation of IL-25 and type 2-associated cytokines in the colon.

### Microbiome reconstitution with minimal live biotherapeutic consortia made of succinate-producing strains increases IL-25, IL-13, and IL-5 in the colon but not in the cecum or ileum

The data from the FMT experiments provides causal evidence of the microbiome in promoting TC-dependent cytokine production in the colon. To demonstrate that this effect can be recapitulated by administration of a limited consortium of bacteria actively producing succinate, we first performed reconstitution experiments where AVNM-pretreated mice were recolonized with either a consortium of three known *in vivo* succinate-producing *Bacteroides* and *Prevotella* species (*Bacteroides thetaiotaomicron*, *Bacteroides vulgatus*, *Prevotella copri*) (Ferreyra *et al*., 2014; Iljazovic *et al*., 2021; Louis and Flint, 2017), a consortium of three common gut bacteria with no known *in vivo* succinate production (*Alistipes shaii*, *Eubacterium eligens*, *Dorea formicigenerans*), or with succinate-producing *Bacteroides* and *Prevotella* species in heat-killed form as control (See Methods). Mice receiving the succinate-producing live bacteria consortium showed significant enrichment in IL-25, IL-13, and IL-5 in the colon (FDR adjusted p-value ANOVA with Tukey post-hoc <0.05) compared to the control treatments (**Fig. 3A**). Mice treated with the non-succinate-producing consortium displayed interleukins levels that were on average higher compared to the treatment with heat-killed bacteria (**Fig. 3A**). This indicates that these strains may produce small amounts of succinate *in vivo* that promotes the production of IL-25, IL-13, and IL-5, or otherwise may alter the existing microbiome to produce more succinate than the heat-killed bacterial treatment. Alternatively, the consortium of bacteria not known to produce succinate may be inducing TC-derived IL-25 production using another and to date unknown signaling pathway. As in the stool transplant experiments, we did not observe significant differences in IL-25 (FDR adjusted p-value ANOVA with Tukey post-hoc > 0.05) in either the cecum or in the ileum of these animals (**Fig. 3 B, C**). We confirmed that the administration of the succinate-producing bacteria consortium significantly increased succinate concentration by performing targeted metabolomics for succinate and other microbiota-related SCFAs from colonic content from untreated mice (PBS), AVNM-treated, AVNM plus vancomycin FMT treated, AVNM plus succinate producing bacteria or AVNM plus non-succinate producers (See Methods). Mice gavaged with the succinate-producing consortium after AVNM treatment displayed significantly higher levels of succinate compared to mice that received AVNM treatment and not producers (two-sample t-test p <0.05) (**Fig. 3D,** **Fig. S5**). No difference in any other SCFAs was observed (**Fig. 3D,** **Fig. S5**) (two-sample t-test FDR-adjusted p >0.05).

**Figure 3:**
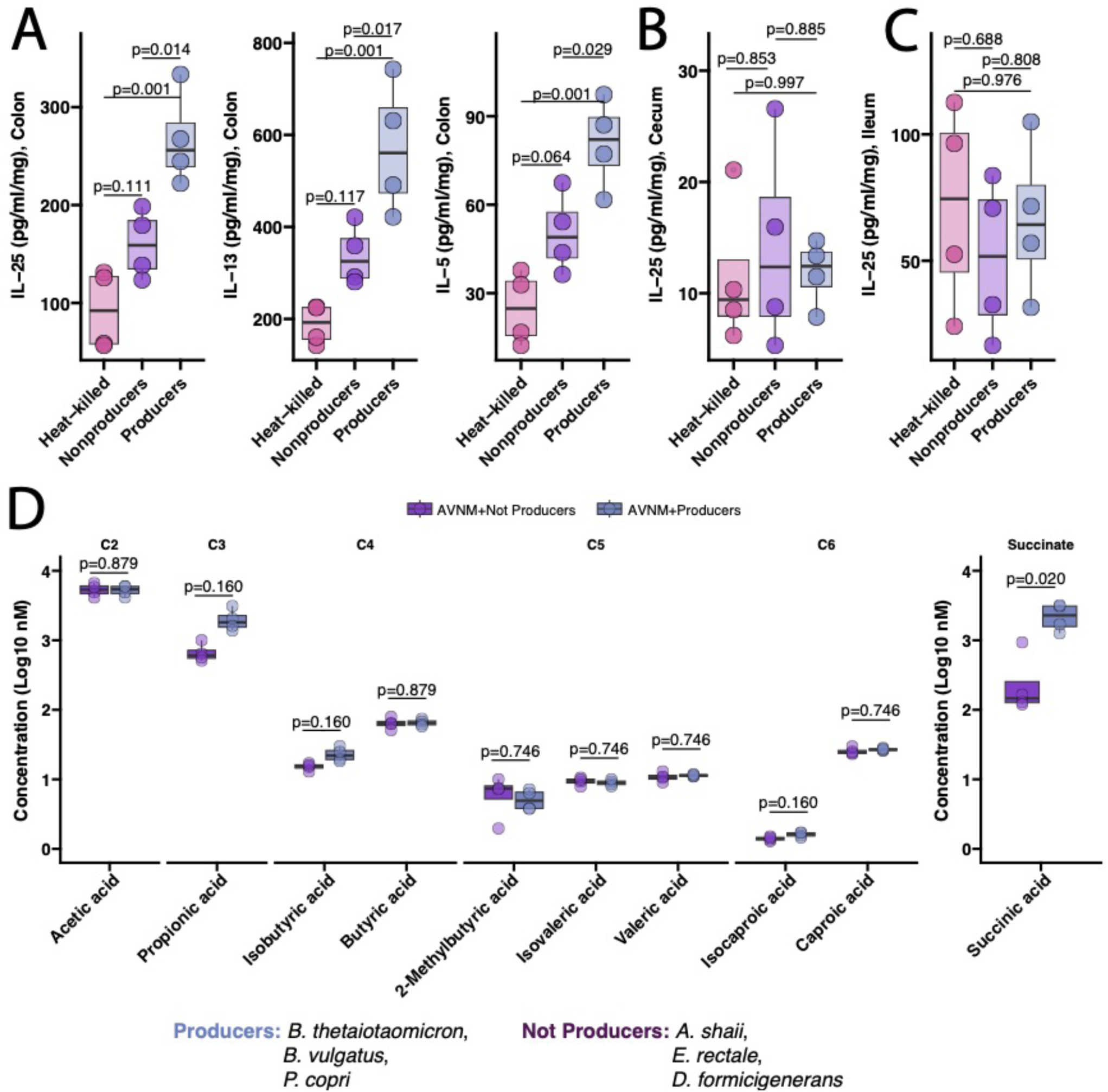
Consortia of succinate-producing bacteria increase Type 2 cytokines in the colon which also corresponds to higher colonic levels of succinate. **A.** IL-25, IL-5, and IL-25 protein levels were measured by ELISA (See Methods) in C57BL/6 wild-type mice receiving succinate-, heat-killed-succinate-, and non-succinate-producing bacterial consortia. **B.** and **C.** IL-25 protein levels were also measured in the cecum and the ileum. **D.** Targeted metabolomics was performed to quantify levels of acetic acid, propionic acid, butyric acid, succinic acid, 2-methylbutryic acid, isolaveric acid, valeric acid, and caproic acid in the proximal colonic luminal contents of C57BL/6 wild-type mice receiving the succinate-or non-succinate-producing bacterial consortia. Protein concentration in the lysate (pg/mL) was normalized by the total protein mass generated in the sample (in mg)

We then sought to confirm that TCs expansion in the proximal colon occurs in response to the succinate produced by these bacteria by comparing the frequency of DCLK1+EPCAM+ cells in mice that were orally gavaged with wild-type *B. thetaiotaomicron*, the succinate production-deficient *B. thetaiotaomicron* fumarate reductase knockout ( frd) (Spiga *et al*., 2017) or heat-killed *B. thetaiotaomicron*. Mice administered with the wild-type, succinate-producing *B. thetaiotaomicron* displayed a significantly higher percentage of DCLK1+EPCAM+ cells compared to those administered with the *B. thetaiotaomicron* frd knockout or the heat-killed *B. thetaiotaomicron* (FDR adjusted p-value ANOVA with Tukey post-hoc <0.05) in the proximal colon (**Fig. 4A**) but not in the ileum (**Fig. 4B**). This also corresponded to higher concentrations of IL-25 mRNA expression (**Fig. 4C**). Interestingly, mice receiving *B. thetaiotaomicron* frd showed higher levels of TCs and IL-25 expression compared to mice gavaged with the heat-killed strain suggesting that TCs may be also responding to other *B. thetaiotaomicron* frd-produced component, or the interaction of *B. thetaiotaomicron* frd with remaining microbial residents. To evaluate the effect that succinate production by *B. thetaiotaomicron* has on other main colonic cytokines (type 1 and type 2) we examined broad cytokine levels in colonic tissues from mice orally administered with wild-type *B. thetaiotaomicron* or with *B. thetaiotaomicron* frd. We found no differences in colonic levels of several type 1 cytokines including (IL-1b, IL-2, IL-21, IL-22) (Student t-Test p >0.05) (**Fig. 4D**), but we found a significant reduction in type 2-associated cytokines levels such as IL-31 and IL-33 in mice administered with *B. thetaiotaomicron* frd (**Fig. 4E**), confirming the dependency of type 2-related cytokines on microbially-produced succinate.

**Figure 4:**
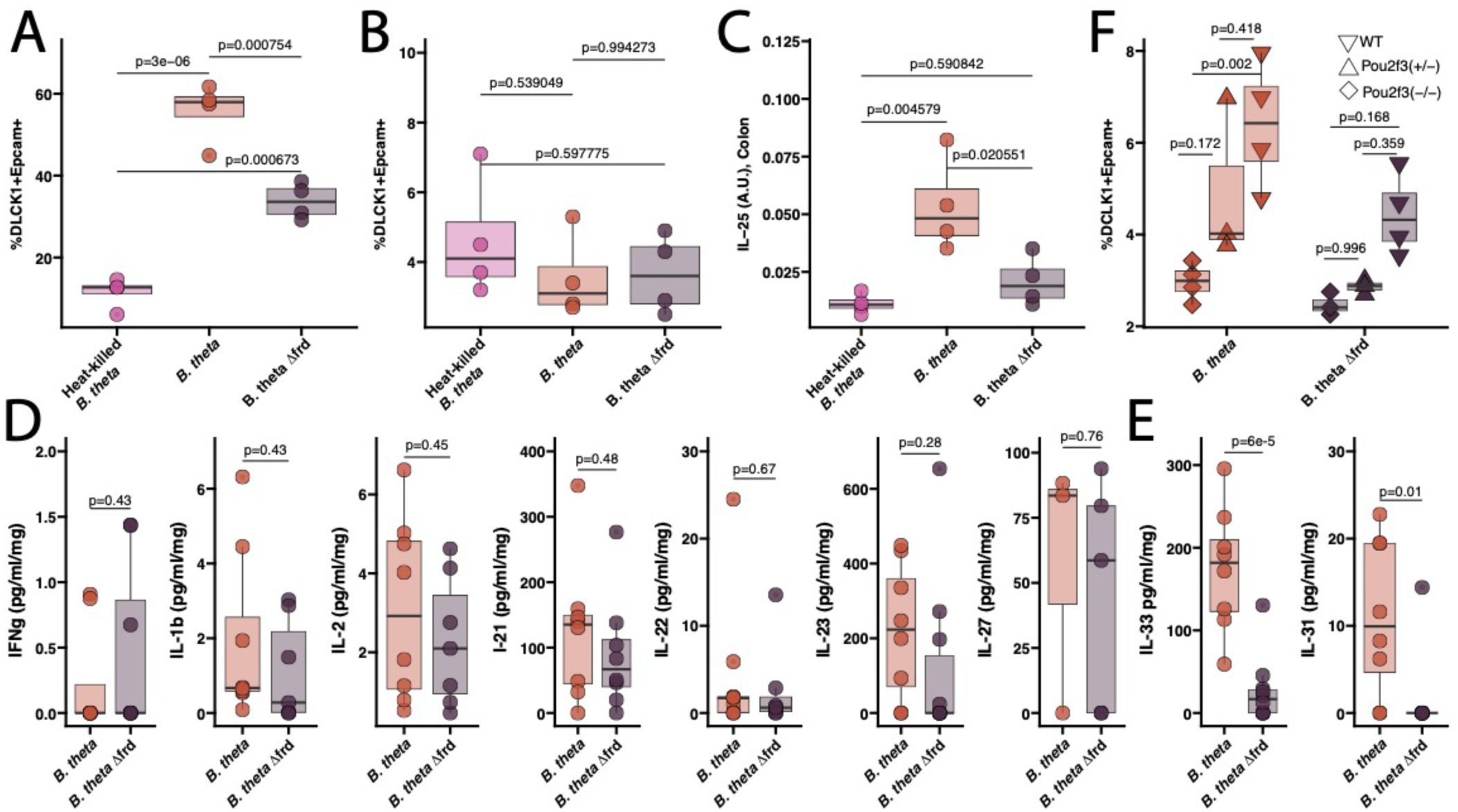
Colonic induction of tuft cells and related cytokines is dependent on the presence of succinate-producing *Bacteroides thetaiotaomicron* and Pou2f3-dependent tuft cells. AVNM-treated mice were orally gavaged with one among heat-killed *Bacteroides thetaiotaomicron*, live *B. thetaiotaomicron*, or *B. thetaiotaomicron* Δfrd. **A.** DCLK1+EPCAM+ cells expressed as a percentage of total cells in the proximal colon of C57BL/6 wild-type mice. **B.** DCLK1+ and EPCAM+ cells expressed as a percentage of total cells in the ileum of C57BL/6 wild-type mice treated with heat-killed *Bacteroides thetaiotaomicron*, live *B. thetaiotaomicron*, or *B. thetaiotaomicron* ΔFRD. **C.** Relative mRNA expression measured by RT-qPCR of IL-25 in the proximal colon of C57BL/6 wild-type mice treated with heat-killed *Bacteroides thetaiotaomicron*, live *B. thetaiotaomicron*, or *B. thetaiotaomicron* Δfrd. **D and E** Type 1 and Type 2-associated cytokines measured by Luminex Multiplex ELISA in the proximal colon of C57BL/6 wild-type mice treated with *B. thetaiotaomicron* or *B. thetaiotaomicron* Δfrd. **F.** DCLK1+EPCAM+ cells expressed as a percentage of total cells in the proximal colon of Pou2f3^+/-^, Pou2f3^-/-^, and C57BL/6 wild-type (WT) mice treated with *B. thetaiotaomicron* or *B. thetaiotaomicron* Δfrd.

To evaluate that TCs expansion in response to succinate produced by *B. thetaiotaomicron* is dependent on the presence of TCs we replicated the experiment where *B. thetaiotaomicron* or *B. thetaiotaomicron* frd were administered to C57BL/6 wild-type, *Pou2f3*^+/-^ or *Pou2f3*^-/-^ mice. The transcription factor *Pou2f3* is crucial for the differentiation of DCLK1+ TCs systemically, including in the GI tract, and its deletion leads to defective mucosal type 2 responses to helminth infection (Gerbe *et al*., 2016). We found that while the TC fraction was significantly higher in C57BL/6 wild-type mice compared to *Pou2f3*^+/-^ and *Pou2f3*^-/-^ when administered with wild-type *B. thetaiotaomicron*, no difference was observed in mice receiving the *B. thetaiotaomicron* frd (p-value for interaction term of linear model <0.05) (**Fig. 4F**). This suggests that the presence of *Pou2f3*-dependent TCs is needed to achieve succinate-dependent DCLK1+ TCs expansion in the colon.

Taken together these data provide evidence that precise microbiome supplementation with succinate-producing bacteria increases succinate concentration and type 2 cytokines in the colon, with no effect on type 1 cytokines. Additionally, these data provide evidence of succinate production by the microbiome as a main driver of TCs expansion and IL-25 accumulation in the colon.

### Prophylactic administration of succinate producing bacteria promotes TCs-mediated protection against *C. difficile*-induced morbidity and mortality

Though succinate accumulation in the lumen promotes *C. difficile* expansion in the colon (Ferreyra *et al*., 2014), it is unknown how this translates into host susceptibility to *C. difficile*-induced disease. Because oral administration of recombinant IL-25 induces eosinophilia that protects mice against *C. difficile* morbidity and mortality with no significant differences in *C. difficile* luminal titer (Buonomo *et al*., 2016), we hypothesized that there is a succinate-centered circuit in the colon by which the microbiome, the host and *C. difficile* interact. Specifically, we posit that microbiome-produced succinate is sensed by TCs to initiate an immune cascade that culminates in protection against *C. difficile*-caused morbidity and mortality.

To test the hypothesis, we leveraged a *C. difficile* infection model (Theriot *et al*., 2011) which we combined with our previously published approach for the adoptive transfer of *C. difficile* disease ameliorating consortia (Buffie *et al*., 2015; Dsouza *et al*., 2022) (See Methods) (**Fig. 5A**). Following AVNM treatment, we administered a suspension containing either the live succinate-producing consortium (*B. thetaiotaomicron*, *B. vulgatus*, *P. copri*), the heat-killed consortium, or PBS to animals prior to infection with *C. difficile* VPI 10463 spores. Adoptive transfer of the consortium alone significantly ameliorated *C. difficile* infection by increasing survival (log-rank survival test p-values = 0.051 and 0.014; succinate producers vs. PBS, and succinate producers vs. heat-killed, respectively) (**Fig. 5B**) and resulted in lower weight loss compared to both controls (two-samples t-test Benjamini-Hochberg adjusted p-value <0.05 comparing weight loss in succinate producers vs. PBS, and succinate producers vs. heat-killed independently at different time points.) (**Fig. 5C**).

**Figure 5:**
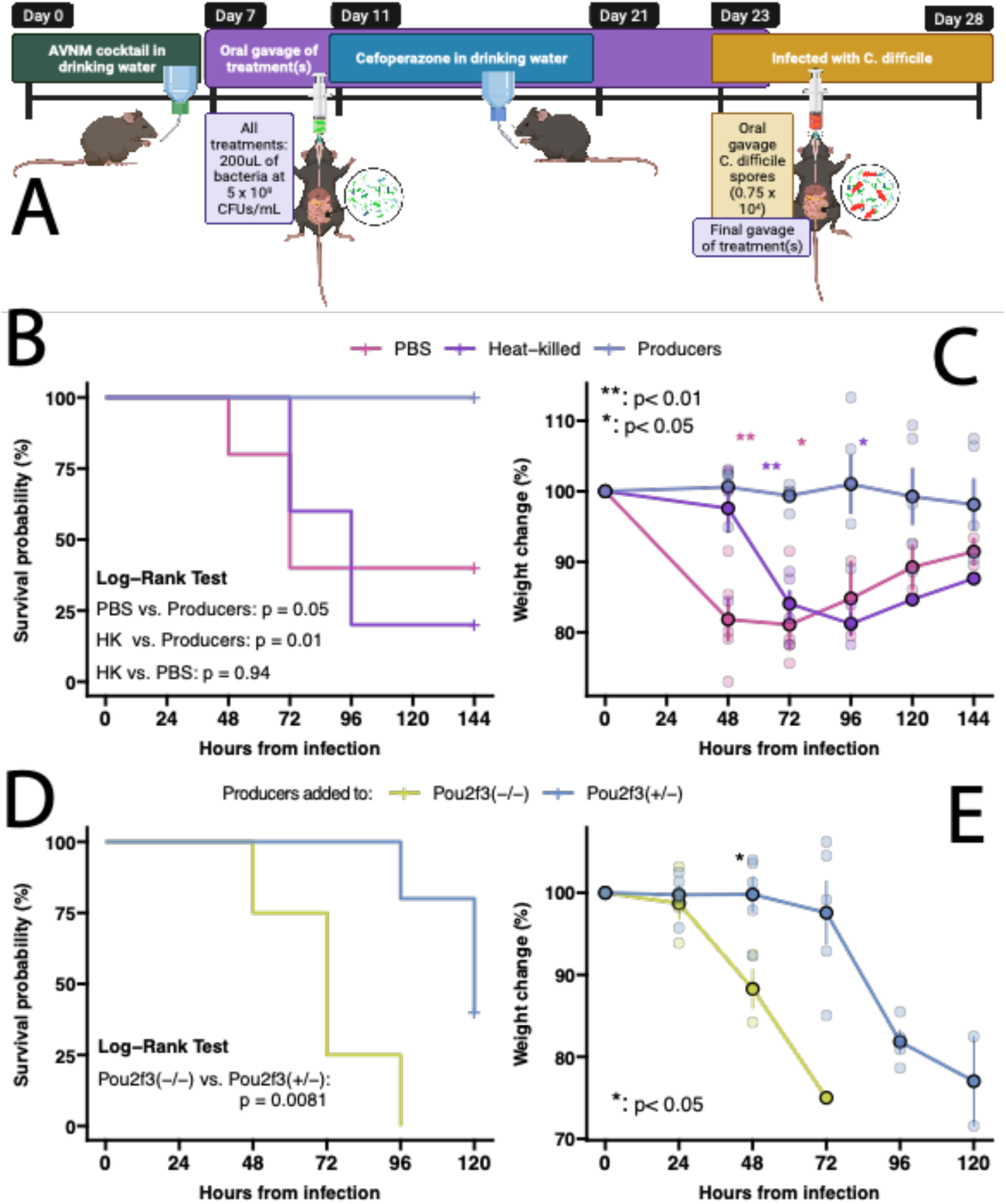
Protection from *C. difficile* morbidity and mortality is obtained by prophylactic administration of succinate-producing live bacteria consortia and depends on the presence of Pou2f3-dependent tuft cells. **A.** Experimental diagram of administration of bacterial consortia and subsequent infection with *Clostridoides difficile*. **B**. Survival of C57BL/6 wild-type mice infected with *C. difficile* following treated with succinate-producing, heat-killed-succinate-producing bacterial consortia, or sterile PBS. **C**. Percentage of weight change in C57BL/6 wild-type mice infected with *C. difficile* following treated with succinate-producing, heat-killed-succinate-producing bacterial consortia, or sterile PBS. **D**. Survival of Pou2f3^(-/-)^ and Pou2f3^(+/-)^ mice infected with *C. difficile* following treated with succinate-producing, heat-killed-succinate-producing bacterial consortia, or sterile PBS. **E**. Percentage of weight change in Pou2f3^(-/-)^ and Pou2f3^(+/-)^ mice infected with *C. difficile* following treated with succinate-, heat-killed-succinate-producing bacterial consortia, or sterile PBS.

To mechanistically prove that the observed protection against *C. difficile* was mediated by the presence of TCs we compared survival and weight loss in *Pou2f3^+^* ^/-^ and *Pou2f3*^-/-^ mice that were adoptively transferred with the consortium of succinate-producing bacteria. As predicted, *Pou2f3^+^* ^/-^ and *Pou2f3*^-/-^ animals displayed increased death and weight loss despite being administered with the succinate producers (**Fig. 5D,E**). However, Pou2f3*^+^* ^/-^ mice displayed significantly higher survival (p value < 0.0013 log-rank survival test) (**Fig. 5D**) and lower weight loss compared to *Pou2f3*^-/-^ (**Fig. 5E**) (two-samples t-test Benjamini-Hochberg adjusted p-value <0.05 comparing weight loss between the two genotypes) confirming the dependency of the protection on the presence of TC in the colon.

Altogether our data provide evidence that the administration of succinate-producing bacteria promotes TC expansion and production of type 2 cytokines to protect against *C. difficile*-induced morbidity and mortality.

## Discussion

Microbiome-produced metabolites modulate innate and adaptive immune responses in the intestine (Geva-Zatorsky *et al*., 2017; Zheng *et al*., 2020). Most mechanistic studies on this topic have focused on unveiling the causal role by which microbial-derived metabolic end-products (such as SCFAs and SBAs) preserve immune homeostasis (Atarashi *et al*., 2013; Atarashi *et al*., 2011; Foley *et al*., 2021; Tanoue *et al*., 2016), and on determining the routes by which gut pathobionts promote inflammation (Atarashi *et al*., 2017; Britton Graham *et al*., 2020).

Succinate is an intermediary metabolite that is produced during the degradation of dietary fibers into other fatty acids (Fernández-Veledo and Vendrell, 2019). Succinate accumulation has often been associated with higher incidence of obesity and IBD (Serena *et al*., 2018), but it has also been found to promote gluconeogenesis and brain signaling (de Vadder and Mithieux, 2018). While succinate accumulation in the SI during invasion by parasites promotes the expansion of TCs with implications for tissue regeneration and parasite expulsion (Ting and von Moltke, 2019; von Moltke *et al*., 2016), it also stimulates growth and activates virulence for different enteric pathogens such as *C. difficile* and *Salmonella enterica*.

We hypothesized that colonic microbial succinate accumulation is a central signal mediating the interaction between the resident microbiota, the host, and enteric pathogens. Specifically, we hypothesized that in the absence of succinate consuming bacteria (e.g., commensal *Clostridia*) succinate accumulation is sensed by colonic TCs to initiate a program allowing the host to temporarily resist intestinal distress and pathogen-induced disease.

In support of this hypothesis, we demonstrate that selective colonic microbiome perturbation leading to the enrichment of succinate-producing microbes (through the selective depletion of succinate consumers) induces TCs expansion and consequent production of type 2 immune cytokines. We show the causality of the phenotype through microbiome reconstitution experiments with stool transplants that are enriched for succinate-producing bacteria, as well as by using defined consortia of known succinate-producing commensals. We demonstrate the dependency of this phenotype on microbial-produced succinate as knocking out succinate production in *B. thetaiotaomicron* leads to significantly lower TC number and type 2 cytokines levels in the colon, while not affecting levels of type 1 cytokines. This effect is localized to the colon; no effect is observed in the small intestine or in the cecum. Furthermore, we show that this phenotype requires live succinate-producing bacteria as heat-killed succinate-producing bacteria do not stimulate this colonic TC-dependent circuit.

Following reports that exogenous administration of IL-25 protects from *C. difficile* induced-colonic damage (Buonomo *et al*., 2016), we show that prophylactic administration of succinate-producing live bacteria protects from *C. difficile* morbidity, and mortality. Importantly, we show that this mechanism is mediated by the presence of TCs as protection is lost in TC knockout mice. This is a significant advancement to studies aimed at exploiting the microbiome to control *C. difficile*-induced disease; our approach provides access to a novel microbiome-regulated axis for immune-mediated *C. difficile* control that can be complementarily explored in addition to current microbiome-based efforts at preventing *C. difficile* disease which are focused on using bacteria intended to directly inhibit this pathogen in the intestine (Bobilev *et al*., 2019; Buffie *et al*., 2015).

At homeostasis, succinate is a metabolic intermediate in the conversion of dietary fibers to health-promoting metabolites including short-chain fatty acids (Fernández-Veledo and Vendrell, 2019). Abnormal accumulation of microbiome-derived succinate in the intestine is a signature of gastrointestinal dysbiosis and is associated with the emergence of different diseases including IBD and obesity (Mills and O’Neill, 2014; Serena *et al*., 2018). Considering this, and the results of this study, we propose that succinate-sensing by colonic TCs is a sentinel mechanism that evolved to temporarily counteract the loss of succinate-to-SCFA converters during dysbiosis which may have a role in containing damage that is caused by dysbiosis-thriving opportunistic pathogens. Perhaps *C. difficile* activates virulence factors in the presence of commensal succinate to overcome the increased intestinal protection provided by TCs in the presence of a succinate-enriched microbiome.

## MATERIALS AND METHODS

### Mice

All animal studies were approved by the UMass Chan Institutional Animal Care and Use Committee (Protocols A-1993-17 and PROTO202100184) in accordance with National Institutes of Health guidelines. All experiments were performed with mice 8–12–weeks of age. C57BL/6J wild-type Specific Pathogen Free (SPF) mice of both sexes were purchased from The Jackson Laboratory (Bar Harbor, ME). C57BL/6J-Pou2f3em1Cbwi/J mice were used to generate *Pou2f*^+/-^ and *Pou2f*^-/-^ animals in-house. Animals were acclimatized to housing facilities for 4 weeks before use in experiments.

### Fecal pellet collection

Mice were placed into separate, autoclaved plastic beakers until 3 fecal pellets were produced. Immediately after production, individual; fecal pellets were transferred using sterile toothpicks into a microfuge tube and flash-frozen in liquid nitrogen.

### Antibiotic administration experiments

The approach follows previous work published by us and others (Foley *et al*., 2021). C57BL6/6J 8-10 week old female SPF mice (n = 6-12 per treatment group, depending on the experiment) were treated with metronidazole (1 g l^-1^), vancomycin (500 mg l^-1^), or an antibiotic cocktail, AVNM (a combination of ampicilllin (1 g l^-1^), vancomycin (500 mg l^-1^), neomycin (1 g l^-1^), and metronidazole (1 g l^-1^)) suspended in phosphate-buffered saline (PBS) or PBS as control. Treatment was performed *via* oral gavage every 12 hours for a total of 7 days. 12 hours after the final antibiotic gavage, mice were sacrificed using carbon dioxide. Tissue samples and intestinal contents were extracted and immediately flash-frozen for immune phenotype quantification. Feces were collected before, during, and after antibiotic treatment and bacterial DNA was extracted as part of the shotgun metagenomic sequencing analysis detailed below.

### Stool matter transplant experiments

8–10-week-old C57BL6/6J female SPF mice (n = 6 per treatment group) classified as recipients were either pre-treated with AVNM in their drinking water for 7 days to deplete the resident microbiome or left on standard acidified drinking water. SPF mice (n=4 per group) classified as donors were pre-treated with vancomycin (500 mg l^-1^) or standard acidified drinking water. After 7 days of antibiotic pre-treatment, all mice were returned to standard acidified drinking water for the remainder of the experiment. 24 hours were allowed to pass between the removal of the antibiotics and the first fecal transplant to allow for antibiotic washout. Donor mice were placed individually in autoclaved plastic beakers until they produced three fecal pellets. Fecal pellets from donor mice in each group were pooled and collected into a 15mL conical tube containing 5mL PBS and resuspended. The fibrous matter was pelleted at 300x g for 5 minutes and removed from the fecal suspension to facilitate passage through the gavage needle. The fecal suspension from either untreated SPF mice or from mice previously treated with vancomycin was orally introduced to recipient mice at 0.2mL/g bodyweight. This was repeated every day for 5 days, with donor feces collected and suspended fresh each day. 24 hours after the final transplant, all mice were sacrificed by carbon dioxide euthanasia. Tissue samples and intestinal content extracts for immune phenotype quantification were collected and flash-frozen. Feces were collected before, during, and after the experiment as described above, and samples of each suspension were taken for bacterial DNA extraction as part of the 16S rRNA gene pyrosequencing analysis detailed below.

### Bacterial growth for live consortia

All bacterial work was performed in a Coy^TM^ anerobic chamber available in the UMass Chan Center for Microbiome Research. All strains were grown in BD Difco™ Reinforced Clostridial Media (BD 218081). All bacterial species were previously determined to have approximately 1x10^8^ colony forming units/mL at an optical density (600nm) of 1 when grown for 48 hours. Bacterial strains were grown in 20mL of media in sterile, anaerobic media bottles at 37C at 50 RPMs for 48 hours. OD600 was determined, and individual strains were pelleted at 10,000 x g for 10 minutes. Bacterial pellets were resuspended in the appropriate volume of anaerobic, sterile PBS to produce 3.33mL of the strain at an OD600 of either 1, or 5. Bacteria were then pooled to produce the consortia into either group A, a consortium of bacteria known to produce succinate *in vivo*, (*Bacteroides thetaiotaomicron* VPI 5482, *Bacteroides vulgatus* NCTC 11154, *Prevotella copri* DSM 18205), or group B, a consortium of bacteria not known to product succinate *in vivo*, (*Alistipes shaii* BAA 1179, *Eubacterium rectale* ATCC 33656, *Dorea formicigenerans* ATCC2 7755), so that the final volume of each consortium was 10mL. This process was repeated daily for each administration of the live consortia. This same approach was used for the experiments comparing immune induction by *Bacteroides thetaiotaomicron* VPI 5482 and the succinate production-deficient *B. thetaiotaomicron* frd (Spiga *et al*., 2017).

### Live consortia administration experiments

8–10-week-old C57BL6/6J female SPF mice (n = 5 per treatment group) were pre-treated with AVNM (ampicillin (1 g l^-1^), vancomycin (500 mg l^-1^), neomycin (1 g l^-1^), and metronidazole (1 g l^-1^)), in their drinking water for 7 days to deplete the resident microbiome (see above). Mice were then administered *via* oral gavage (as above) to either (1) group A, “succinate-producers” described above, (2) group B, “non-producers” described above, or (3) the bacteria from group A after heat-killing for 5 minutes at 65°C at an OD600 of 1. Mice were gavaged at 0.2mL/g bodyweight. The administration was repeated every 24 hours for 4 days. 24 hours after the last administration of the live consortia, mice were euthanized by carbon dioxide. Tissue samples and intestinal extracts were collected and flash-frozen. Feces were collected before, during, and after the experiment as described above for bacterial DNA extraction as part of the 16S rRNA gene pyrosequencing detailed below. This same experimental protocol was used in the assays comparing phenotype induction by *Bacteroides thetaiotaomicron* VPI 5482 or the succinate production-deficient *B. thetaiotaomicron* frd (Spiga *et al*., 2017) in wild-type C57BL6/6J, Pou2f3^+/-^ and Pou2f3^-/-^, 8–10-week-old female SPF mice.

### Clostridioides difficile infection experiments

We evaluated the response of mice receiving different live bacterial consortia to *C. difficile* infection by adapting the animal model first described in (Theriot *et al*., 2011). Briefly, 8–10-week-old C57BL6/6J female SPF mice were pre-treated with AVNM for one week. After 1 day of AVNM washout, mice were orally gavaged once a day by the “succinate-producers” (see above) or heat-killed producers at an OD600 of 5 (approximately 5x10^8^ CFUs/mL), or PBS daily for 14 days at 0.2mL/g bodyweight. Three days from the start of bacterial administration mice were administered cefoperazone (0.5 mg/ml) (MP Bioworks, cat# 199695) in sterile drinking water for 10 days. After 2 days to let cefoperazone wash out, mice were then orally gavaged with 10^5^ CFUs of *C. difficile* strain VPI 10463 (ATCC 43255). Animals were assessed for symptoms such as inappetence (lack of appetite), diarrhea, and hunching. Animals were euthanized if they lost 20% of their initial baseline weight or exhibited any severe clinical signs listed above. Similar *C. difficile* infection experiments were performed where succinate-producing bacteria were orally gavaged into Pou2f3^+/-^ and Pou2f3^-/-^, 8–10-week-old female SPF mice.

### Tissue preparation and cytokine measurement

Cecal tissue was flushed with sterile PBS and sectioned into 0.5cm sections before flash freezing. Ileal and proximal colon tissue were manually evacuated and sectioned into 0.5cm sections before flash freezing. Protein lysates from intestinal tissue were generated as described in (Foley *et al*., 2021) (i.e., benchtop homogenization in tubes containing Lysing Matrix D (MP Biomedical) beads and lysis buffer (20 mM Tris pH 7.4, 120 mM NaCl, 1 mM EDTA, 1% Triton-X-100, 0.5% sodium deoxycholate, 1× protease inhibitor cocktail [Roche])). Cytokine protein levels were measured by ELISA (IL-17E, IL-5, and IL-13 Duo-Set, R&D Systems). Cytokine levels were normalized to total protein concentration in the tissue lysates using the DC Protein Assay (BioRad). RNA from epithelial cells and tissue was isolated by following the TRIzol extraction manufacture protocol. Total RNA was used for RT–PCR. Complementary DNA was generated using iScript Reverse Transcription Supermix (Invitrogen, catalog no. 18080-044). For RT–qPCR, cDNA was mixed with appropriate primers (Supplementary Table 3) and SYBR green master mix (BioRad, catalog no. 1708882) and run on a Thermocycler T100 (BioRad). Proximal colon lysates were used to measure levels of IL-1b, IL-2, IL-21, IL-22, IL-31 and IL-33 cytokines with multiplexed-ELISA assay with Luminex 200 Multiplex Bio-Plex 200 System (EMD Millipore, Billerica, MA, USA) using a Milliplex Map kit (EMD Millipore).

### Succinate and Short Chain Fatty Acids measurement

Quantitation of C2 to C6 short-chain fatty acids (SCFAs) and succinic acid was carried out as previously described in (Han *et al*., 2015) by the Uvic-Genome BC Proteomics Centre. Briefly, serially diluted standard solutions of SCFAs and succinic acid were prepared with the use of their standard substances in 60% acetonitrile. An internal standard (IS) solution of the isotope-labeled version of SCFAs and succinic acid was prepared using 13C6-3-nitrophenylhydrazine and following the derivatizing procedure described in the publication. The samples were precisely weighed into 2-mL homogenizing tubes. 60% acetonitrile at 10 μL per mg of raw material was added. The samples were homogenized on a MM 400 mill mixer with the aid of two metal beads at 30 Hz for 3 min, followed by centrifugal clarification at 21,000 rpm and 5 °C for 10 min. 20 μL of the clear supernatant of each sample or standard solution was mixed in turn with 80 μL of 200-mM 3-nitrophenylhydrazine solution and 80 μL of 150-mM EDC-6% pyridine solution. The mixtures were incubated at 40 °C for 30 min. After the reaction, each solution was diluted 10-fold with the IS solution. 10-μL aliquots of the resultant solutions were injected into a C18 (2.1*150 mm, 1.8 μm) column to run LC-MRM/MS on a Waters UPLC system coupled to a Sciex 4000 QTRAP mass spectrometer with negative-ion detection. The mobile phase was 0.01% formic acid in water (A) and 0.01% formic acid in acetonitrile (B) for binary-solvent gradient elution of 15% to 90% B over 15 min, at 40 °C and 0.35 mL/min. Linear-regression calibration curves were constructed with the data acquired from injections of the standard solutions. Concentrations of the detected analytes in the samples were calculated by interpolating the calibration curves with the peak area ratios measured from injections of the sample solutions.

### Flow cytometry

Mouse intestines were opened longitudinally and vortexed in a 50-ml conical tube containing Hanks’ balanced salt solution supplemented with 5% heat-inactivated FBS and 10 mM HEPES, pH 7.2. Epithelial cells were isolated by rotating the tissues in a pre-digestion medium (RPMI medium, 5% heat-inactivated FBS, 10 mM HEPES, pH 7.2, and 10 mM EDTA) for 30 min at 37 °. Cells were stained with antibodies for CD326 (BioLegend, catalog no. G8.8, 1:200), CD45.2 (BioLegend, catalog no. 30-F11), anti-DCLK1 (Abcam; ab31704) and LIVE/DEAD.

### 16s rRNA sequencing and bioinformatics

The bacterial 16S rRNA gene (variable regions V3 to V4) was subjected to PCR amplification using the universal 341F and 806R barcoded primers for Illumina sequencing. Using the SequalPrep Normalization kit, the products were pooled into sequencing libraries in equimolar amounts and sequenced on the Illumina MiSeq platform using v3 chemistry for 2 × 300 bp reads. The forward and reverse amplicon sequencing reads were dereplicated and sequences were inferred using dada2 (Callahan *et al*., 2016) as in (Wipperman *et al*., 2021).

### Statistical analysis of host phenotypes

We conducted several statistical analyses to investigate variations in cytokine protein levels, gene expression, tuft cell numbers, metabolite levels, mouse survival in response to *C. difficile* infection, and the effect of treatment on weight loss.

To assess differences in cytokine protein levels, gene expression, and tuft cell numbers, we employed a two-step analysis. Initially, we conducted an Analysis of Variance (ANOVA) followed by Tukey post-hoc tests to compare multiple groups. In the case of metabolite levels for mice administered with *B. thetaiotaomicron* or *B. thetaiotaomicron* Δfrd, we used two-sample t-tests. We determined significant associations at a False Discovery Rate (FDR) value of 0.05, ensuring that the observed differences were statistically reliable.

For evaluating differences in mouse survival following *C. difficile* infection due to different treatments or mouse genotypes, we employed log-rank tests. To investigate the impact of treatment on weight loss after C. difficile infection, we run Benjamini-Hochberg-corrected two-samples t-test at different time points as in (Dsouza *et al*., 2022). All statistical analyses were carried out using the R programming language.

### Statistical and machine learning modeling of 16S rRNA data

Differences in the abundance of DADA2-identified amplicon sequencing variants (ASVs) were evaluated using DESeq2 (Love *et al*., 2014) in “R”. Prediction of colonic IL-25 as a function of the abundance of 16S rRNA-determined ASVs was performed by building and running Random Forest Regression (RFR) models as in (Wipperman *et al*., 2021). Permutated Variable Importance and Accumulated Local Effect algorithms were used to determine the significance and directionality of the modeling-identified microbial abundances-IL-25 relationships. Results from the RFR analysis were confirmed by running Elastic Net and Bayesian Variable Selection linear regression models using in house code, see (Bucci *et al*., 2016). Significant associations were determined at a False Discovery Rate value of 0.05.

## Acknowledgments

We would like to acknowledge Dr. Alexander Rudensky and Dr. Michael Buchert for insightful conversation. We would like to thank Dr. Sebastian Winter for providing the *B. thetaiotaomicron* frd strain.

## Author Contributions

Conceptualization: VB, AR, BAM, TDK

Laboratory experiments: TDK, SC, ALZ, SF, BMM

Microbiome sequencing and informatics: TDK, BM, DVW, SKB

Data analysis and interpretation: TDK, SC, VB, AR

Funding acquisition: VB, BAM, AR

Supervision: VB, AR

Writing – original draft: TDK, VB, AR

Writing – review and editing: all authors

## Data and materials availability

All data associated with this study are in the paper or supplementary materials. Microbiome sequencing data are being deposited in the SRA with an accession number provided upon paper acceptance. Code to perform all the reported analysis is available in Zenodo upon paper acceptance.

AR and VB are co-corresponding authors. Co-authorship and author order were determined by the recognition that the integration of the nuances of microbiome data, mathematical modeling, and immunology are different skill sets found in different laboratory environments. Each was an important component of the validity and message of this manuscript.

## Supplementary Figures

**Figure S1:**
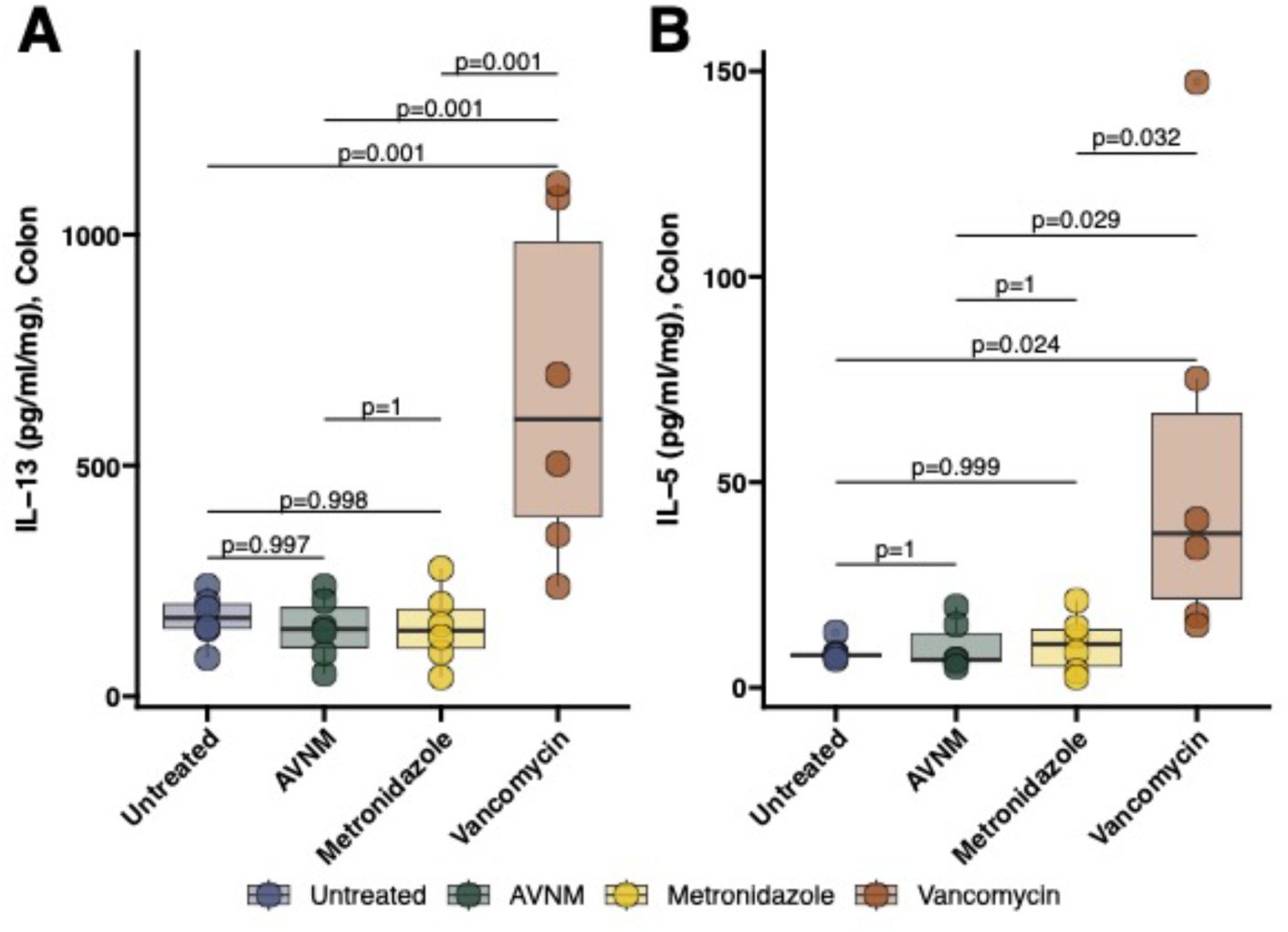
Vancomycin administration increases the concentration of interleukins IL-5 and IL-13 in the proximal colon. **A.** IL-5 and **B.** IL-13 were measured by ELISA in the proximal colon of WT C57BL/6 mice treated with different antibiotics, or left untreated. Protein concentration in the tissue lysate (pg/mL) was normalized by the total protein mass generated in the sample (in mg).

**Figure S2:**
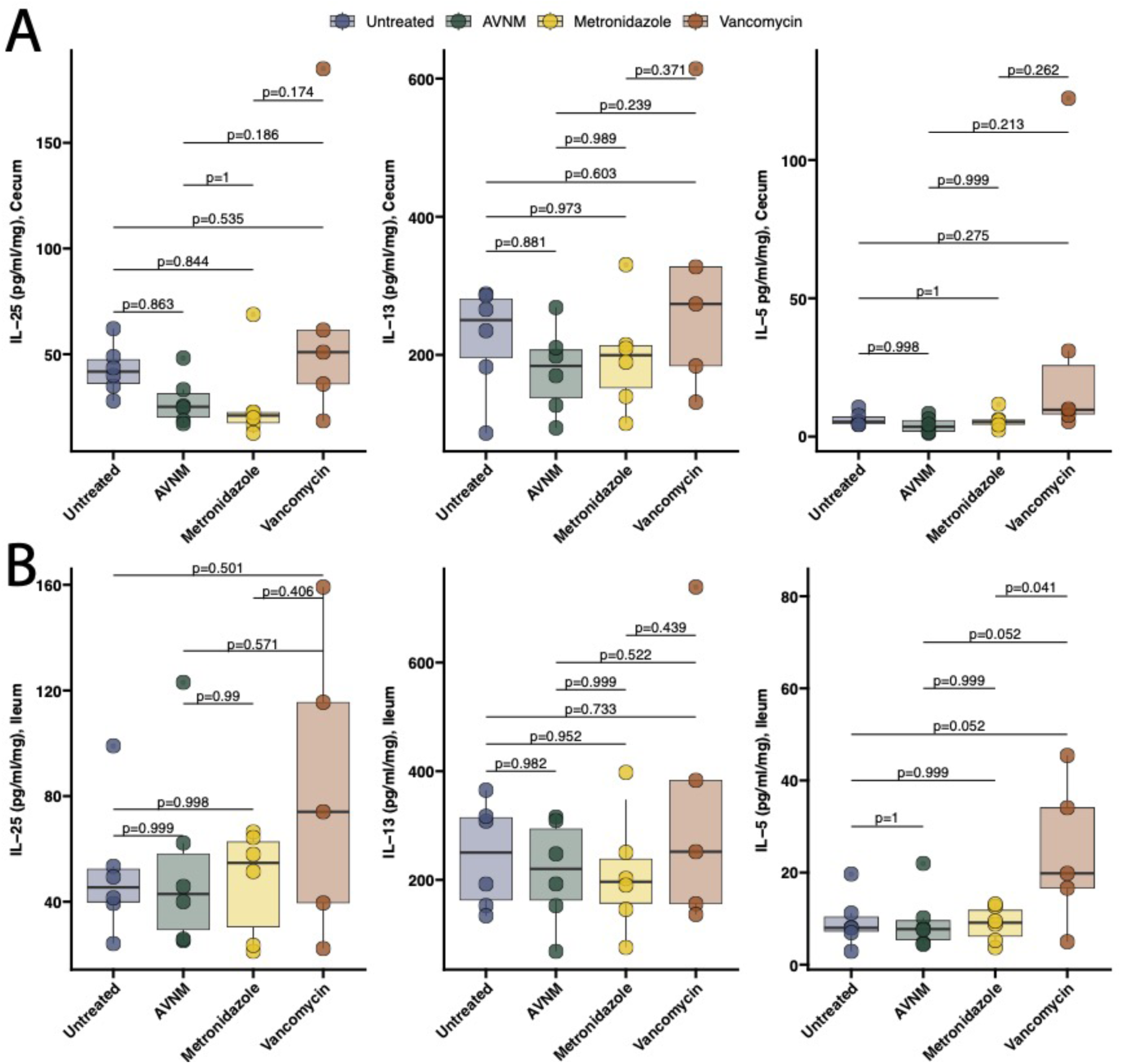
Vancomycin administration *does not* significantly increase IL-25, IL-5, or IL-13 in the cecum or ileum. IL-25, IL-13, and IL-5 were measured with ELISA in the cecum **A.** or in the Ileum **B.** of WT C57BL/6 mice treated with different antibiotics, or left untreated. Protein concentration in the tissue lysate (pg/mL) was normalized by the total protein mass generated in the sample (in mg).

**Figure S3:**
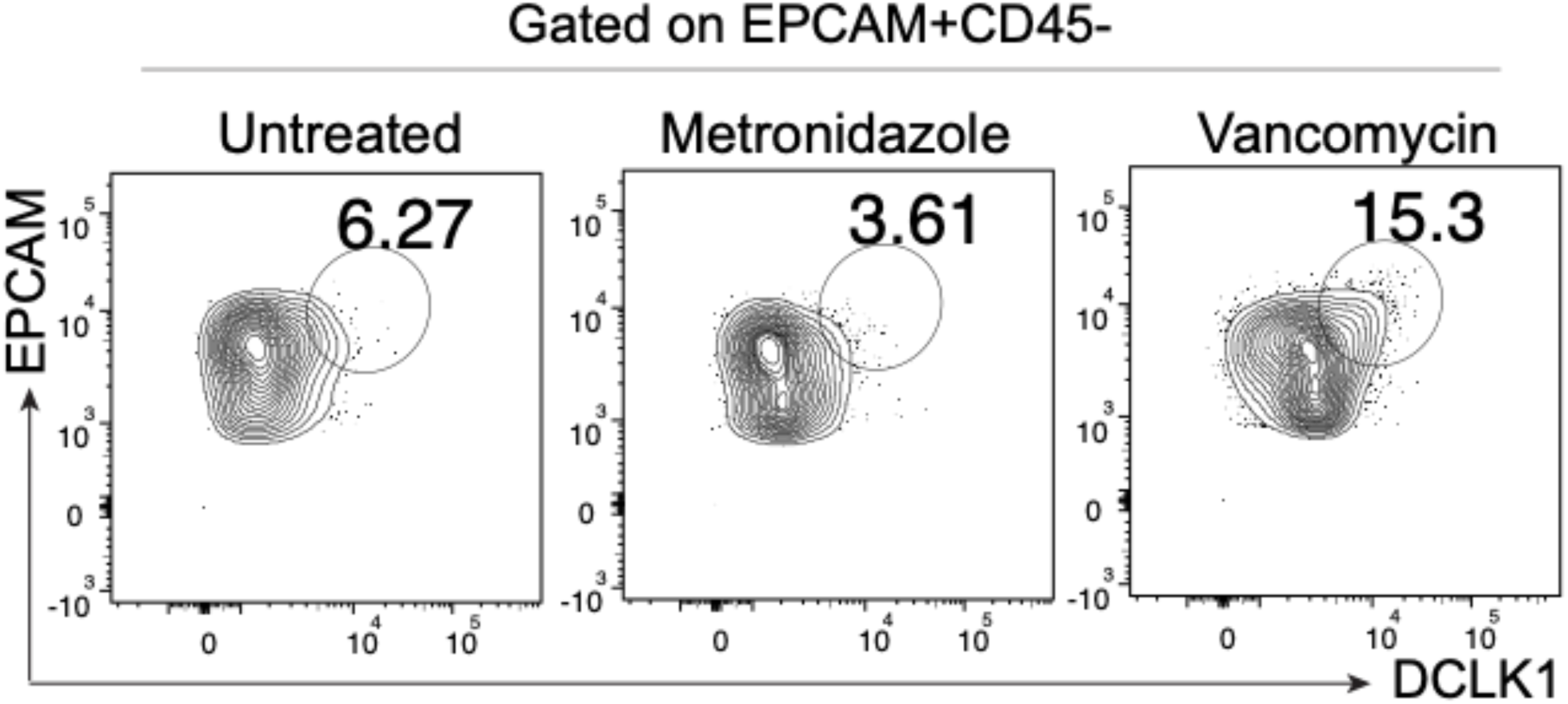
Representative gating of DCLK1+ Epcam+ CD45-tuft cells *via* flow cytometry. Flow cytometry was used to estimate the percentage of DCLK1+ EPCAM+ CD45-cells compared to total cells in C57BL/6 wild-type mice treated with various antibiotics or left untreated. Mice treated with vancomycin had a significantly higher percentage of DCLK1+ Epcam+ CD45-tuft cells than mice treated with metronidazole or left untreated.

**Figure S4:**
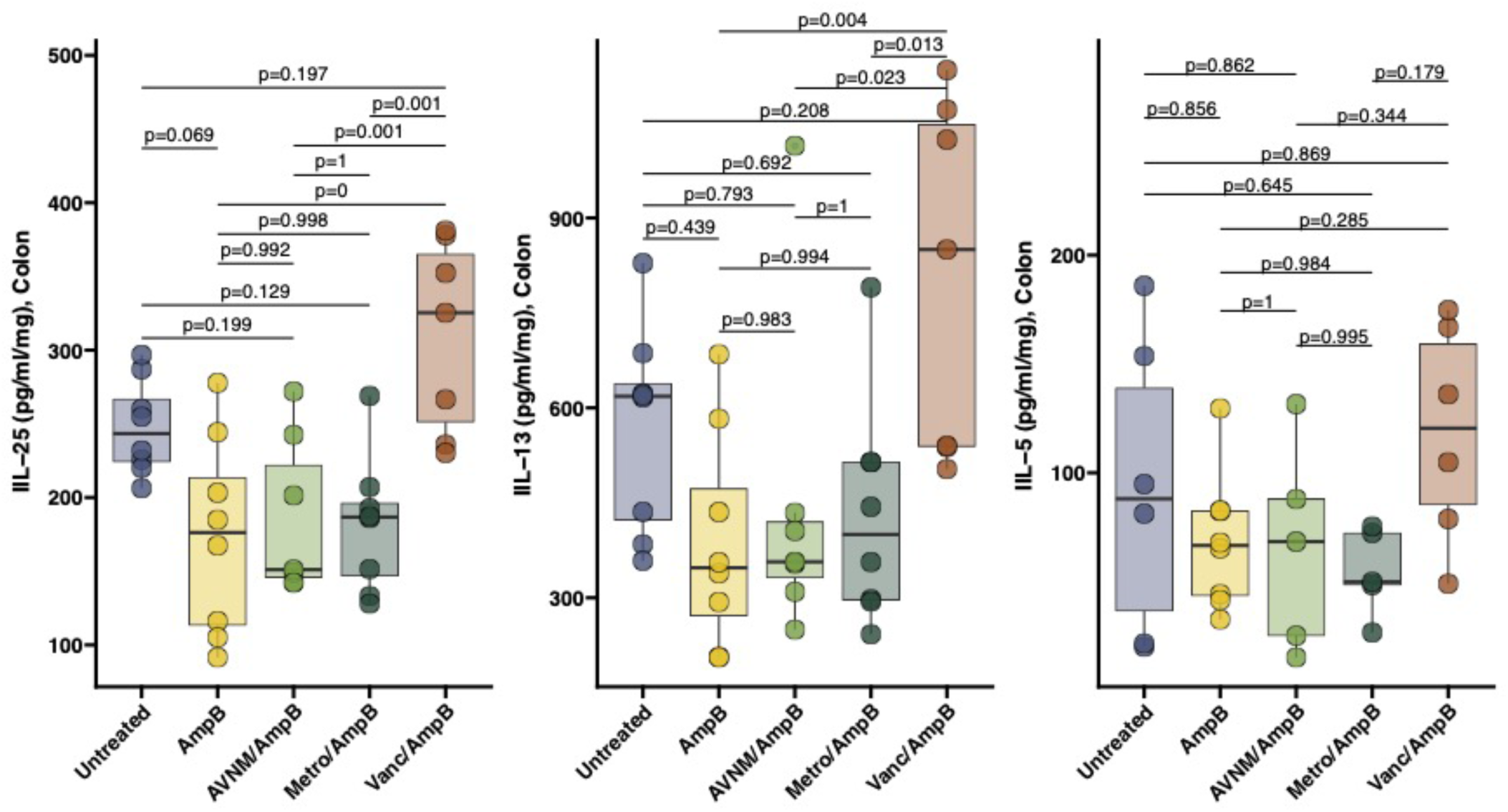
Amphotericin B administration does not change the ability of vancomycin administration to upregulate IL-25, IL5, and IL-13 in the proximal colon. **(A)** IL-25 measured by ELISA in the proximal colon of C57BL/6 wild-type (WT) mice treated with antibiotics in combination with amphotericin B (ampB). **(B)** IL-13 measured by ELISA in the proximal colon of WT mice treated with antibiotics in combination with ampB. **(C)** IL-5 measured by ELISA in the proximal colon of WT mice treated with antibiotics in combination with ampB. Protein concentration in the tissue lysate (pg/mL) was normalized by the total protein mass generated in the sample (in mg).

**Figure S5:**
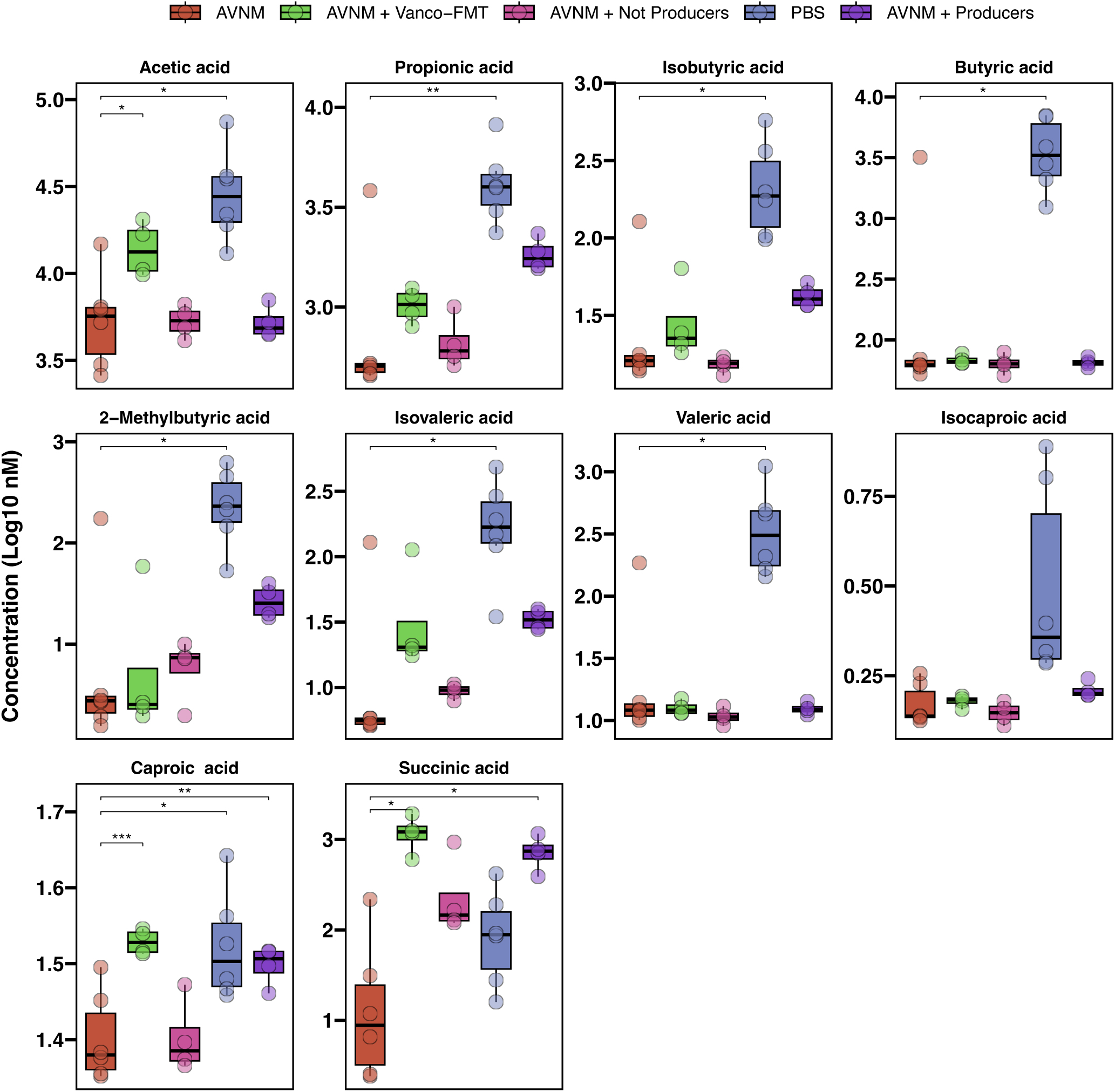
Targeted metabolomics for succinate and SCFAs in mice treated with AVNM (baseline), AVNM + vancomycin FMT, AVNM + succinate-producing bacteria, AVNM + non-succinate producing bacteria, or untreated (PBS). ANOVA with Tukey post hoc was run to determine significant differences compared to AVNM (baseline). *** p < 0.0001, ** p < 0.01, * p < 0.05

## References

1. Akdis, C.A., P.D. Arkwright, M.C. Brüggen, W. Busse, M. Gadina, E. Guttman-Yassky, K. Kabashima, Y. Mitamura, L. Vian, J. Wu, and O. Palomares. 2020. Type 2 immunity in the skin and lungs. Allergy 75:1582–1605.

2. Aoki, R., A. Aoki-Yoshida, C. Suzuki, and Y. Takayama. 2018. Indole-3-Pyruvic Acid, an Aryl Hydrocarbon Receptor Activator, Suppresses Experimental Colitis in Mice. J Immunol 201:3683–3693.

3. Arpaia, N., C. Campbell, X. Fan, S. Dikiy, J. van der Veeken, P. deRoos, H. Liu, J.R. Cross, K. Pfeffer, P.J. Coffer, and A.Y. Rudensky. 2013. Metabolites produced by commensal bacteria promote peripheral regulatory T-cell generation. Nature 504:451–455.

4. Atarashi, K., W. Suda, C. Luo, T. Kawaguchi, I. Motoo, S. Narushima, Y. Kiguchi, K. Yasuma, E. Watanabe, T. Tanoue, C.A. Thaiss, M. Sato, K. Toyooka, H.S. Said, H. Yamagami, S.A. Rice, D. Gevers, R.C. Johnson, J.A. Segre, K. Chen, J.K. Kolls, E. Elinav, H. Morita, R.J. Xavier, M. Hattori, and K. Honda. 2017. Ectopic colonization of oral bacteria in the intestine drives T(H)1 cell induction and inflammation. Science (New York, N.Y.) 358:359–365.

5. Atarashi, K., T. Tanoue, K. Oshima, W. Suda, Y. Nagano, H. Nishikawa, S. Fukuda, T. Saito, S. Narushima, K. Hase, S. Kim, J.V. Fritz, P. Wilmes, S. Ueha, K. Matsushima, H. Ohno, B. Olle, S. Sakaguchi, T. Taniguchi, H. Morita, M. Hattori, and K. Honda. 2013. Treg induction by a rationally selected mixture of Clostridia strains from the human microbiota. Nature 500:232–236.

6. Atarashi, K., T. Tanoue, T. Shima, A. Imaoka, T. Kuwahara, Y. Momose, G. Cheng, S. Yamasaki, T. Saito, Y. Ohba, T. Taniguchi, K. Takeda, S. Hori, Ivanov, II, Y. Umesaki, K. Itoh, and K. Honda. 2011. Induction of colonic regulatory T cells by indigenous Clostridium species. Science 331:337–341.

7. Banerjee, A., C.A. Herring, B. Chen, H. Kim, A.J. Simmons, A.N. Southard-Smith, M.M. Allaman, J.R. White, M.C. Macedonia, E.T. McKinley, M.A. Ramirez-Solano, E.A. Scoville, Q. Liu, K.T. Wilson, R.J. Coffey, M.K. Washington, J.A. Goettel, and K.S. Lau. 2020. Succinate Produced by Intestinal Microbes Promotes Specification of Tuft Cells to Suppress Ileal Inflammation. Gastroenterology 159:2101–2115.e2105.

8. Begum, N., A. Harzandi, S. Lee, M. Uhlen, D.L. Moyes, and S. Shoaie. 2022. Host-mycobiome metabolic interactions in health and disease. Gut Microbes 14:2121576.

9. Blander, J.M., R.S. Longman, I.D. Iliev, G.F. Sonnenberg, and D. Artis. 2017. Regulation of inflammation by microbiota interactions with the host. Nature Immunology 18:851–860.

10. Bobilev, D., S. Bhattarai, R. Menon, B. Klein, S. Reddy, B. Olle, B. Roberts, V. Bucci, and J. Norman. 2019. 1953. VE303, a Rationally Designed Bacterial Consortium for Prevention of Recurrent Clostridioides difficile (C. Difficile) infection (rCDI), Stably Restores the Gut Microbiota After Vancomycin (vanco)-Induced Dysbiosis in Adult Healthy Volunteers (HV). Open Forum Infect Dis 6:S60-S60.

11. Britton Graham, J., J. Contijoch Eduardo, P. Spindler Matthew, V. Aggarwala, B. Dogan, G. Bongers, L. San Mateo, A. Baltus, A. Das, D. Gevers, J. Borody Thomas, O. Kaakoush Nadeem, A. Kamm Michael, H. Mitchell, S. Paramsothy, C. Clemente Jose, J.-F. Colombel, W. Simpson Kenneth, C. Dubinsky Marla, A. Grinspan, and J. Faith Jeremiah. 2020. Defined microbiota transplant restores Th17/RORγt+ regulatory T cell balance in mice colonized with inflammatory bowel disease microbiotas. Proceedings of the National Academy of Sciences 117:21536–21545.

12. Bucci, V., B. Tzen, N. Li, M. Simmons, T. Tanoue, E. Bogart, L. Deng, V. Yeliseyev, M.L. Delaney, Q. Liu, B. Olle, R.R. Stein, K. Honda, L. Bry, and G.K. Gerber. 2016. MDSINE: Microbial Dynamical Systems INference Engine for microbiome time-series analyses. Genome Biol 17:121.

13. Buffie, C.G., V. Bucci, R.R. Stein, P.T. McKenney, L. Ling, A. Gobourne, D. No, H. Liu, M. Kinnebrew, A. Viale, E. Littmann, M.R.M. van den Brink, R.R. Jenq, Y. Taur, C. Sander, J.R. Cross, N.C. Toussaint, J.B. Xavier, and E.G. Pamer. 2015. Precision microbiome reconstitution restores bile acid mediated resistance to Clostridium difficile. Nature 517:205–208.

14. Buonomo, E.L., C.A. Cowardin, M.G. Wilson, M.M. Saleh, P. Pramoonjago, and W.A. Petri, Jr. 2016. Microbiota-Regulated IL-25 Increases Eosinophil Number to Provide Protection during Clostridium difficile Infection. Cell Rep 16:432–443.

15. Callahan, B.J., P.J. McMurdie, M.J. Rosen, A.W. Han, A.J.A. Johnson, and S.P. Holmes. 2016. DADA2: High-resolution sample inference from Illumina amplicon data. Nature Methods 13:581–583.

16. Catlett, J.L., J. Catazaro, M. Cashman, S. Carr, R. Powers, M.B. Cohen, and N.R. Buan. 2020. Metabolic Feedback Inhibition Influences Metabolite Secretion by the Human Gut Symbiont Bacteroides thetaiotaomicron. mSystems 5:

17. Chen, J., Y. Wang, L. Shen, Y. Xiu, and B. Wang. 2022. Could IL-25 be a potential therapeutic target for intestinal inflammatory diseases? Cytokine & Growth Factor Reviews

18. Cox, J.R., S.M. Cruickshank, and A.E. Saunders. 2021. Maintenance of Barrier Tissue Integrity by Unconventional Lymphocytes. Front Immunol 12:670471.

19. de Vadder, F., and G. Mithieux. 2018. Gut-brain signaling in energy homeostasis: the unexpected role of microbiota-derived succinate. Journal of Endocrinology 236:R105–R108.

20. Dsouza, M., R. Menon, E. Crossette, S.K. Bhattarai, J. Schneider, Y.-G. Kim, S. Reddy, S. Caballero, C. Felix, L. Cornacchione, J. Hendrickson, A.R. Watson, S.S. Minot, N. Greenfield, L. Schopf, R. Szabady, J. Patarroyo, W. Smith, P. Harrison, E.J. Kuijper, C.P. Kelly, B. Olle, D. Bobilev, J.L. Silber, V. Bucci, B. Roberts, J. Faith, and J.M. Norman. 2022. Colonization of the live biotherapeutic product VE303 and modulation of the microbiota and metabolites in healthy volunteers. Cell Host & Microbe 30:583–598.e588.

21. Fernández-Veledo, S., and J. Vendrell. 2019. Gut microbiota-derived succinate: Friend or foe in human metabolic diseases? Reviews in Endocrine and Metabolic Disorders 20:439–447.

22. Ferreyra, J.A., K.J. Wu, A.J. Hryckowian, D.M. Bouley, B.C. Weimer, and J.L. Sonnenburg. 2014. Gut microbiota-produced succinate promotes C. difficile infection after antibiotic treatment or motility disturbance. Cell Host Microbe 16:770–777.

23. Foley Sage, E., J. Dente Michael, X. Lei, F. Sallis Benjamin, B. Loew Ethan, M. Meza-Segura, A. Fitzgerald Katherine, and A. McCormick Beth. 2022. Microbial Metabolites Orchestrate a Distinct Multi-Tiered Regulatory Network in the Intestinal Epithelium That Directs P-Glycoprotein Expression. mBio 13:e01993-01922.

24. Foley, S.E., C. Tuohy, M. Dunford, M.J. Grey, H. De Luca, C. Cawley, R.L. Szabady, A. Maldonado-Contreras, J.M. Houghton, D.V. Ward, R.J. Mrsny, and B.A. McCormick. 2021. Gut microbiota regulation of P-glycoprotein in the intestinal epithelium in maintenance of homeostasis. Microbiome 9:183.

25. Fremder, M., S.W. Kim, A. Khamaysi, L. Shimshilashvili, H. Eini-Rider, I.S. Park, U. Hadad, J.H. Cheon, and E. Ohana. 2021. A transepithelial pathway delivers succinate to macrophages, thus perpetuating their pro-inflammatory metabolic state. Cell Reports 36:109521.

26. Gerbe, F., E. Sidot, D.J. Smyth, M. Ohmoto, I. Matsumoto, V. Dardalhon, P. Cesses, L. Garnier, M. Pouzolles, B. Brulin, M. Bruschi, Y. Harcus, V.S. Zimmermann, N. Taylor, R.M. Maizels, and P. Jay. 2016. Intestinal epithelial tuft cells initiate type 2 mucosal immunity to helminth parasites. Nature 529:226–230.

27. Geva-Zatorsky, N., E. Sefik, L. Kua, L. Pasman, T.G. Tan, A. Ortiz-Lopez, T.B. Yanortsang, L. Yang, R. Jupp, D. Mathis, C. Benoist, and D.L. Kasper. 2017. Mining the Human Gut Microbiota for Immunomodulatory Organisms. Cell 168:928–943.e911.

28. Gieseck, R.L., 3rd, M.S. Wilson, and T.A. Wynn. 2018. Type 2 immunity in tissue repair and fibrosis. Nat Rev Immunol 18:62-76.

29. Goettel, J.A., R. Gandhi, J.E. Kenison, A. Yeste, G. Murugaiyan, S. Sambanthamoorthy, A.E. Griffith, B. Patel, D.S. Shouval, H.L. Weiner, S.B. Snapper, and F.J. Quintana. 2016. AHR Activation Is Protective against Colitis Driven by T Cells in Humanized Mice. Cell Rep 17:1318–1329.

30. Han, J., K. Lin, C. Sequeira, and C.H. Borchers. 2015. An isotope-labeled chemical derivatization method for the quantitation of short-chain fatty acids in human feces by liquid chromatography-tandem mass spectrometry. Anal Chim Acta 854:86–94.

31. Howitt, M.R., S. Lavoie, M. Michaud, A.M. Blum, S.V. Tran, J.V. Weinstock, C.A. Gallini, K. Redding, R.F. Margolskee, L.C. Osborne, D. Artis, and W.S. Garrett. 2016. Tuft cells, taste-chemosensory cells, orchestrate parasite type 2 immunity in the gut. Science 351:1329–1333.

32. Ikeyama, N., T. Murakami, A. Toyoda, H. Mori, T. Iino, M. Ohkuma, and M. Sakamoto. 2020. Microbial interaction between the succinate-utilizing bacterium Phascolarctobacterium faecium and the gut commensal Bacteroides thetaiotaomicron. Microbiologyopen 9:e1111.

33. Iljazovic, A., U. Roy, E.J.C. Gálvez, T.R. Lesker, B. Zhao, A. Gronow, L. Amend, S.E. Will, J.D. Hofmann, M.C. Pils, K. Schmidt-Hohagen, M. Neumann-Schaal, and T. Strowig. 2021. Perturbation of the gut microbiome by Prevotella spp. enhances host susceptibility to mucosal inflammation. Mucosal Immunology 14:113–124.

34. Isaac, S., J.U. Scher, A. Djukovic, N. Jiménez, D.R. Littman, S.B. Abramson, E.G. Pamer, and C. Ubeda. 2017. Short-and long-term effects of oral vancomycin on the human intestinal microbiota. J Antimicrob Chemother 72:128–136.

35. Kennedy, M.S., and E.B. Chang. 2020. The microbiome: Composition and locations. Prog Mol Biol Transl Sci 176:1–42.

36. Kim, S.H., L.L. Mamuad, D.W. Kim, S.K. Kim, and S.S. Lee. 2016. Fumarate Reductase-Producing Enterococci Reduce Methane Production in Rumen Fermentation In Vitro. J Microbiol Biotechnol 26:558–566.

37. Lei, W., W. Ren, M. Ohmoto, J.F. Urban, Jr., I. Matsumoto, R.F. Margolskee, and P. Jiang. 2018. Activation of intestinal tuft cell-expressed Sucnr1 triggers type 2 immunity in the mouse small intestine. Proc Natl Acad Sci U S A 115:5552–5557.

38. Loke, P., and K. Cadwell. 2018. Getting a Taste for Parasites in the Gut. Immunity 49:16–18.

39. Louis, P., and H.J. Flint. 2017. Formation of propionate and butyrate by the human colonic microbiota. Environmental Microbiology 19:29–41.

40. Love, M.I., W. Huber, and S. Anders. 2014. Moderated estimation of fold change and dispersion for RNA-seq data with DESeq2. Genome Biology 15:550.

41. Luo, X.C., Z.H. Chen, J.B. Xue, D.X. Zhao, C. Lu, Y.H. Li, S.M. Li, Y.W. Du, Q. Liu, P. Wang, M. Liu, and L. Huang. 2019. Infection by the parasitic helminth Trichinella spiralis activates a Tas2r-mediated signaling pathway in intestinal tuft cells. Proc Natl Acad Sci U S A 116:5564–5569.

43. McCarville, J.L., G.Y. Chen, V.D. Cuevas, K. Troha, and J.S. Ayres. 2020. Microbiota Metabolites in Health and Disease. Annu Rev Immunol 38:147–170.

44. Miller, C.N., I. Proekt, J. von Moltke, K.L. Wells, A.R. Rajpurkar, H. Wang, K. Rattay, I.S. Khan, T.C. Metzger, J.L. Pollack, A.C. Fries, W.W. Lwin, E.J. Wigton, A.V. Parent, B. Kyewski, D.J. Erle, K.A. Hogquist, L.M. Steinmetz, R.M. Locksley, and M.S. Anderson. 2018. Thymic tuft cells promote an IL-4-enriched medulla and shape thymocyte development. Nature 559:627–631.

45. Mills, E., and L.A. O’Neill. 2014. Succinate: a metabolic signal in inflammation. Trends Cell Biol 24:313–320.

46. Nadjsombati, M.S., J.W. McGinty, M.R. Lyons-Cohen, J.B. Jaffe, L. DiPeso, C. Schneider, C.N. Miller, J.L. Pollack, G.A. Nagana Gowda, M.F. Fontana, D.J. Erle, M.S. Anderson, R.M. Locksley, D. Raftery, and J. von Moltke. 2018. Detection of Succinate by Intestinal Tuft Cells Triggers a Type 2 Innate Immune Circuit. Immunity 49:33–41 e37.

47. O’Leary, C.E., C. Schneider, and R.M. Locksley. 2019. Tuft Cells—Systemically Dispersed Sensory Epithelia Integrating Immune and Neural Circuitry. Annual Review of Immunology 37:47–72.

48. Qu, D., N. Weygant, R. May, P. Chandrakesan, M. Madhoun, N. Ali, S.M. Sureban, G. An, M.J. Schlosser, and C.W. Houchen. 2015. Ablation of Doublecortin-Like Kinase 1 in the Colonic Epithelium Exacerbates Dextran Sulfate Sodium-Induced Colitis. PLoS One 10:e0134212.

49. Ridlon, J.M., D.J. Kang, P.B. Hylemon, and J.S. Bajaj. 2014. Bile acids and the gut microbiome. Curr Opin Gastroenterol 30:332–338.

50. Schneider, C., C.E. O’Leary, J. von Moltke, H.E. Liang, Q.Y. Ang, P.J. Turnbaugh, S. Radhakrishnan, M. Pellizzon, A. Ma, and R.M. Locksley. 2018a. A Metabolite-Triggered Tuft Cell-ILC2 Circuit Drives Small Intestinal Remodeling. Cell 174:271–284 e214.

51. Schneider, C., C.E. O’Leary, J. von Moltke, H.E. Liang, Q.Y. Ang, P.J. Turnbaugh, S. Radhakrishnan, M. Pellizzon, A. Ma, and R.M. Locksley. 2018b. A Metabolite-Triggered Tuft Cell-ILC2 Circuit Drives Small Intestinal Remodeling. Cell 174:271–284.e214.

52. Schulthess, J., S. Pandey, M. Capitani, K.C. Rue-Albrecht, I. Arnold, F. Franchini, A. Chomka, N.E. Ilott, D.G.W. Johnston, E. Pires, J. McCullagh, S.N. Sansom, C.V. Arancibia-Cárcamo, H.H. Uhlig, and F. Powrie. 2019. The Short Chain Fatty Acid Butyrate Imprints an Antimicrobial Program in Macrophages. Immunity 50:432–445.e437.

53. Serena, C., V. Ceperuelo-Mallafré, N. Keiran, M.I. Queipo-Ortuño, R. Bernal, R. Gomez-Huelgas, M. Urpi-Sarda, M. Sabater, V. Pérez-Brocal, C. Andrés-Lacueva, A. Moya, F.J. Tinahones, J.M. Fernández-Real, J. Vendrell, and S. Fernández-Veledo. 2018. Elevated circulating levels of succinate in human obesity are linked to specific gut microbiota. The ISME Journal 12:1642–1657.

54. Smillie, C.S., J. Sauk, D. Gevers, J. Friedman, J. Sung, I. Youngster, E.L. Hohmann, C. Staley, A. Khoruts, M.J. Sadowsky, J.R. Allegretti, M.B. Smith, R.J. Xavier, and E.J. Alm. 2018. Strain Tracking Reveals the Determinants of Bacterial Engraftment in the Human Gut Following Fecal Microbiota Transplantation. Cell host & microbe 23:229-240.e225.

55. Spiga, L., M.G. Winter, T. Furtado de Carvalho, W. Zhu, E.R. Hughes, C.C. Gillis, C.L. Behrendt, J. Kim, D. Chessa, H.L. Andrews-Polymenis, D.P. Beiting, R.L. Santos, L.V. Hooper, and S.E. Winter. 2017. An Oxidative Central Metabolism Enables Salmonella to Utilize Microbiota-Derived Succinate. Cell Host Microbe 22:291–301.e296.

56. Su, J., T. Chen, X.Y. Ji, C. Liu, P.K. Yadav, R. Wu, P. Yang, and Z. Liu. 2013. IL-25 downregulates Th1/Th17 immune response in an IL-10-dependent manner in inflammatory bowel disease. Inflamm Bowel Dis 19:720–728.

57. Suez, J., N. Zmora, G. Zilberman-Schapira, U. Mor, M. Dori-Bachash, S. Bashiardes, M. Zur, D. Regev-Lehavi, R. Ben-Zeev Brik, S. Federici, M. Horn, Y. Cohen, A.E. Moor, D. Zeevi, T. Korem, E. Kotler, A. Harmelin, S. Itzkovitz, N. Maharshak, O. Shibolet, M. Pevsner-Fischer, H. Shapiro, I. Sharon, Z. Halpern, E. Segal, and E. Elinav. 2018. Post-Antibiotic Gut Mucosal Microbiome Reconstitution Is Impaired by Probiotics and Improved by Autologous FMT. Cell 174:1406–1423.e1416.

58. Tanoue, T., K. Atarashi, and K. Honda. 2016. Development and maintenance of intestinal regulatory T cells. Nature Reviews Immunology 16:295–309.

59. Theriot, C.M., C.C. Koumpouras, P.E. Carlson, Bergin, II, D.M. Aronoff, and V.B. Young. 2011. Cefoperazone-treated mice as an experimental platform to assess differential virulence of Clostridium difficile strains. Gut Microbes 2:326–334.

60. Ting, H.-A., and J. von Moltke. 2019. The Immune Function of Tuft Cells at Gut Mucosal Surfaces and Beyond. The Journal of Immunology 202:1321.

61. Tulstrup, M.V., E.G. Christensen, V. Carvalho, C. Linninge, S. Ahrné, O. Højberg, T.R. Licht, and M.I. Bahl. 2015. Antibiotic Treatment Affects Intestinal Permeability and Gut Microbial Composition in Wistar Rats Dependent on Antibiotic Class. PLoS One 10:e0144854.

62. Ubeda, C., V. Bucci, S. Caballero, A. Djukovic, N.C. Toussaint, M. Equinda, L. Lipuma, L. Ling, A. Gobourne, D. No, Y. Taur, R.R. Jenq, M.R. van den Brink, J.B. Xavier, and E.G. Pamer. 2013. Intestinal microbiota containing Barnesiella species cures vancomycin-resistant Enterococcus faecium colonization. Infect Immun 81:965–973.

63. von Moltke, J., M. Ji, H.E. Liang, and R.M. Locksley. 2016. Tuft-cell-derived IL-25 regulates an intestinal ILC2-epithelial response circuit. Nature 529:221–225.

64. Wipperman, M.F., S.K. Bhattarai, C.K. Vorkas, V.S. Maringati, Y. Taur, L. Mathurin, K. McAulay, S.C. Vilbrun, D. Francois, J. Bean, K.F. Walsh, C. Nathan, D.W. Fitzgerald, M.S. Glickman, and V. Bucci. 2021. Gastrointestinal microbiota composition predicts peripheral inflammatory state during treatment of human tuberculosis. Nat Commun 12:1141.

65. Zheng, D., T. Liwinski, and E. Elinav. 2020. Interaction between microbiota and immunity in health and disease. Cell Research 30:492–506.

